# Human DUX4 and mouse Dux interact with STAT1 and broadly inhibit interferon-stimulated gene induction

**DOI:** 10.1101/2022.08.09.503314

**Authors:** Amy E. Spens, Nicholas A. Sutliff, Sean R. Bennett, Amy E. Campbell, Stephen J. Tapscott

**Affiliations:** Human Biology Division, Fred Hutchinson Cancer Research Center, Seattle, WA 98109; Department of Biochemistry and Molecular Genetics, University of Colorado Anschutz Medical Campus, Aurora, CO 80045; Clinical Research Division, Fred Hutchinson Cancer Research Center, Seattle, WA 98109; Department of Neurology, University of Washington, Seattle, WA 98105

**Keywords:** DUX4, DUX, DUXC, STAT1, interferon gamma, interferon beta, interferon stimulated genes

## Abstract

DUX4 activates the first wave of zygotic gene expression in the early embryo. Mis-expression of DUX4 in skeletal muscle causes facioscapulohumeral dystrophy (FSHD), whereas expression in cancers suppresses IFNγ-induction of MHC Class I and contributes to immune evasion. We show that the DUX4 protein interacts with STAT1 and broadly suppresses expression of IFNγ stimulated genes by decreasing bound STAT1 and Pol-II recruitment. Transcriptional suppression of ISGs requires conserved (L)LxxL(L) motifs in the carboxyterminal region of DUX4 and phosphorylation of STAT1 Y701 enhances interaction with DUX4. Consistent with these findings, expression of endogenous DUX4 in FSHD muscle cells and the CIC-DUX4 fusion containing the DUX4 CTD in a sarcoma cell line inhibit IFNγ-induction of ISGs. Mouse Dux similarly interacted with STAT1 and suppressed IFNγ induction of ISGs. These findings identify an evolved role of the DUXC family in modulating immune signaling pathways with implications for development, cancers, and FSHD.

## INTRODUCTION

Double homeobox (DUX) genes encode a family of transcription factors that originated in placental mammals, consisting of DUXA, DUXB and DUXC subfamilies that all have similar paired homeodomains. The DUXC family is characterized by a small conserved region at the carboxy-terminus of the protein that includes two (L)LxxL(L) motifs and surrounding conserved amino acids (Leidenroth & Hewitt, 2010). Members of this family, including mouse *Dux* and human *DUX4*, are expressed in a brief burst at early stages of development and regulate an initial wave of zygotic gene activation (De Iaco et al., 2017; Hendrickson et al., 2017; Whiddon et al., 2017). While *DUX4* expression has also been reported in testes and thymus (Das & Chadwick, 2016; Snider et al., 2010), it is silenced in most somatic tissues.

Mis-expression of *DUX4* in skeletal muscle is the cause of facioscapulohumeral muscular dystrophy (FSHD) (Campbell et al., 2018; Tawil et al., 2014), the third most prevalent human muscular dystrophy. DUX4 expression in skeletal muscle activates the early embryonic totipotent program, suppresses the skeletal muscle program, and ultimately results in muscle cell loss. Many of the genes induced by DUX4 in skeletal muscle encode proteins that are normally restricted to immune-privileged tissues (Geng et al., 2012) and their expression in skeletal muscle could induce an immune response. In this context it is interesting that FSHD muscle pathology is characterized by focal immune cell infiltrates. However, our prior studies have also suggested that DUX4 might suppress antigen presentation and aspects of an immune response. Expression of DUX4 in cultured muscle cells blocked lenti-viral induction of innate immune response genes such as *IFIH1* (Geng et al., 2012). More recently we reported that expression of DUX4 in primary cancers and engineered cancer cell lines blocks the interferon-gamma (IFNγ) mediated induction of MHC Class I antigen presentation and promotes resistance to immune checkpoint blockade treatments, such as anti-CTLA-4 and anti-PD-1 therapies (Chew et al., 2019). The scope and mechanism(s) of how DUX4 suppresses immune signaling remains unknown.

DUX4 contains one LxxLL and one LLxxL motif at its C-terminal end that are among the most highly conserved regions of DUXC-family (Leidenroth & Hewitt, 2010). LxxLL motifs are alpha-helical protein-interaction domains that were first identified in nuclear-receptor signaling pathways (Heery et al., 1997). Proteins containing LxxLL motifs, such as the <u>P</u>rotein <u>I</u>nhibitor of <u>A</u>ctivated <u>S</u>TAT or PIAS family, have been shown to modulate immune signaling of STATs, IRFs, NF-kB, and other transcription factors (Shuai & Liu, 2005). PIAS proteins block the function of these transcription factors in four ways: preventing DNA binding, recruiting co-repressors, stimulating SUMOylation, or sequestering them within nuclear or sub-nuclear structures (Shuai & Liu, 2005).

In this study we show that a transcriptionally inactive C-terminal fragment of DUX4 is sufficient to block IFNγ-induction of most interferon stimulated genes (ISGs) and this requires the (L)LxxL(L) domains. Immunoprecipitation and mass spectrometry identified the IFNγ-signaling effector STAT1 and several other proteins involved in immune signaling as proteins that interact with the DUX4 C-terminal domain (DUX4-CTD). We show that the DUX4-CTD interacts with STAT1 phosphorylated at Y701 and interferes with stable DNA binding, recruitment of Pol-II, and transcriptional activation of interferon stimulated genes (ISGs). Consistent with these mechanistic studies, endogenous DUX4 in FSHD muscle cells and the CIC-DUX4 fusion protein expressed in a subset of EWSR1-negative small blue round cell sarcomas suppress IFNγ-induction of ISGs. The comparable CTD of mouse Dux containing (L)LxxL(L) motifs similarly interacts with STAT1 and blocks IFNγ stimulation of ISGs. These findings suggest an evolved role of the DUXC family in modulating immune signaling pathways and have implications for the role of DUX4 in development, cancers, and FSHD.

## RESULTS

### DUX4 broadly suppresses interferon-stimulated gene (ISG) induction

Our prior studies showed that DUX4 inhibited ISG induction in response to lentiviral infection and suppressed induction of MHC Class I proteins in response to interferon-gamma (IFNγ, Type-II interferon) (Chew et al., 2019; Geng et al., 2012). To determine whether DUX4 broadly inhibited ISG induction by IFNγ we used the MB135-iDUX4 cell line, a human skeletal muscle cell line with an integrated doxycycline inducible DUX4 (iDUX4) transgene (Jagannathan et al., 2016). (See **Suppl Fig S1A and B** for schematics and sequences of the transgenes used in this study.) Doxycycline induction of DUX4 expression in the MB135-iDUX4 cell line has been validated as an accurate cell model of the transcriptional consequences of DUX4 expression in FSHD muscle cells (Jagannathan et al., 2016) and in the early embryo (Hendrickson et al., 2017; Whiddon et al., 2017). Using a stringent 8-fold induction cut-off (log2 fold-change > 3), RNA-seq showed that IFNγ treatment induced 113 genes, whereas the expression of DUX4 suppressed ISG induction by IFNγ more than 4-fold for 76 (67%) of these genes and more than 2-fold for 102 genes (90%) (**Suppl Table S1**).

Informed by the RNA-seq results, we used RT-qPCR to measure the response of four ISGs that represent different components of the response to immune signaling: the RNA helicase *IFIH1*; the interferon-stimulated exonuclease *ISG20*; the chemoattractant *CXCL9*; and the major histocompatibility complex class II (MHC-II) chaperone *CD74*. IFNγ-induction of all four genes was robustly blocked by DUX4 expression while a DUX4-target gene *ZSCAN4* was strongly induced, indicating that the ISG suppression did not represent a universal block to gene induction (**Fig 1A**, MB135-iDUX4 and **Suppl Fig S2A (**for this and subsequent constructs, **Suppl Fig S2** shows RT-qPCR data from additional independent cell lines together with protein expression and nuclear localization)); whereas doxycycline treatment in the absence of iDUX4 did not suppress ISG induction (**Fig 1A**, MB135 parental). In contrast to DUX4, a paralog in the DUX family, DUXB, did not suppress ISG induction by IFNγ (**Fig 1A**, MB135-iDUXB).

**Figure 1.**
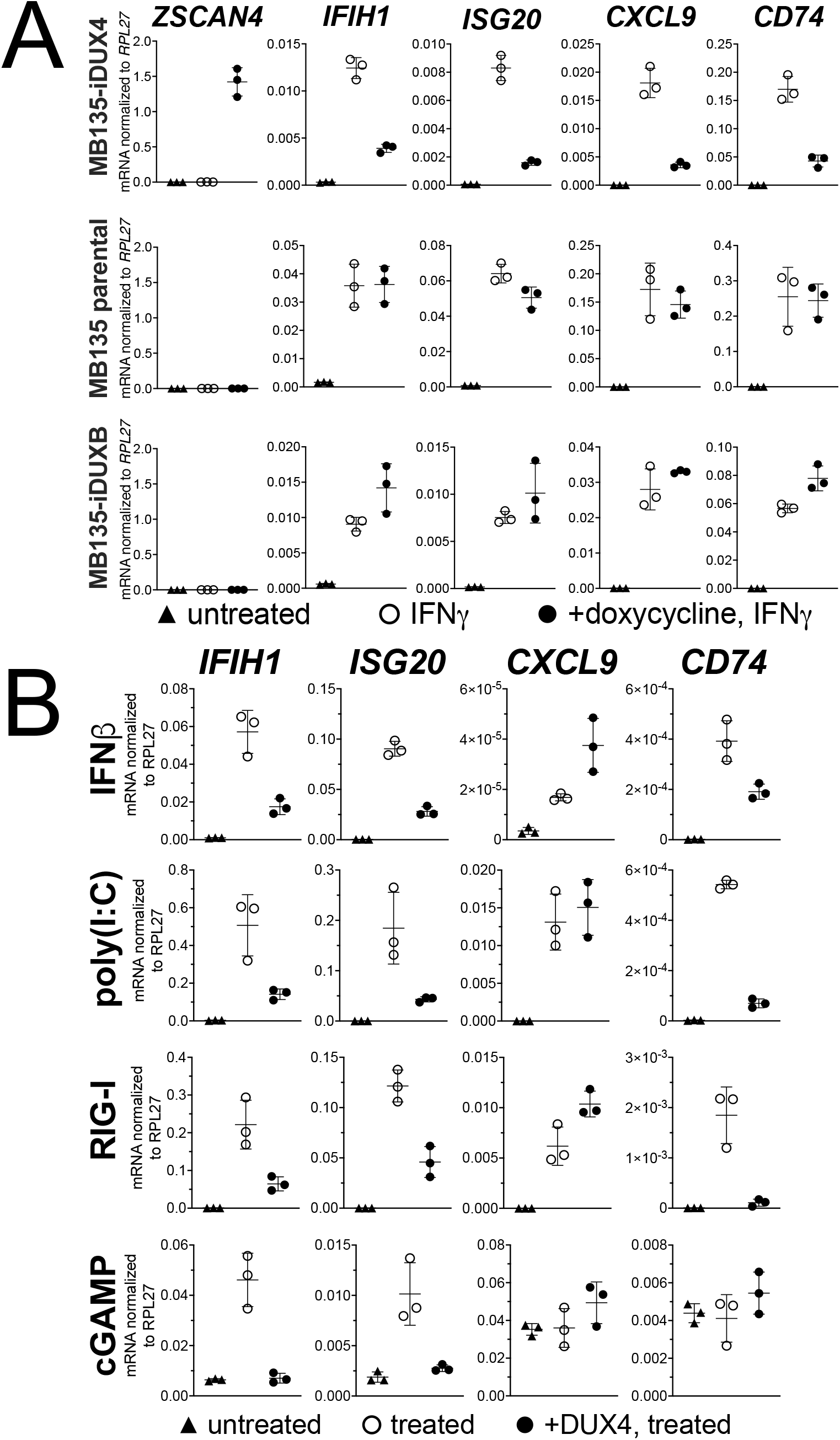
DUX4 suppresses interferon-stimulated gene (ISG) induction. (**A**) MB135 cells expressing doxycycline-inducible DUX4 (MB135-iDUX4), parental MB135 cells, or MB135 cells expressing doxycycline-inducible DUXB (MB135-iDUXB) were untreated, treated with IFNγ, or treated with doxycycline and IFNγ. RT-qPCR was used to evaluate expression of a DUX4 target gene, *ZSCAN4*, and interferon-stimulated genes *IFIH1, ISG20, CXCL9*, and *CD74*. Ct values were normalized to the housekeeping gene *RPL27*. Data represent the mean ±SD of three biological replicates with three technical replicates each. See **Suppl Fig S2** for biological replicates in independent cell lines. (**B**) MB135-iDUX4 cells were untreated, treated with either IFNβ (Type-1 IFN pathway), poly(I:C) (IFIH1/MDA5 pathway), RIG-I ligand (DDX58/RIG-I pathway), or cGAMP (cGAS/STING pathway), or treated with doxycycline and the same immune reagent. RT-qPCR was used to evaluate expression of *IFIH1, ISG20, CXCL9*, and *CD74*. Ct values were normalized to the housekeeping gene *RPL27*. Data represent the mean ±SD of three biological replicates with three technical replicates each.

To determine whether DUX4 also inhibits ISG induction by other innate immune signaling pathways, we transfected the MB135-iDUX4 cells with three different innate immune stimuli: poly(I:C), a long dsRNA mimic to stimulate IFIH1 (MDA5); RIG-I ligand, a short dsRNA with a 5’-ppp to stimulate DDX58 (RIG-I); or cGAMP, a signaling component of the cGAS dsDNA sensing pathway. Additionally, we stimulated the cells with interferon-beta (IFNβ, Type-I interferon), which primarily signals through JAK-STAT pathways via a STAT1-STAT2-IRF9 complex, as opposed to the STAT1 homodimers induced by IFNγ. For all signaling pathways, DUX4 suppressed the induction of a subset of the panel of ISG genes induced by each ligand (**Fig 1B**). One exception, *CXCL9* was induced by IFNβ, poly(I:C), and the RIG-I ligand but not suppressed by DUX4. cGAMP did not induce *CXCL9* or *CD74*, precluding evaluation of the role of DUX4 in regulating these ISGs. These results indicate that DUX4 can modulate the activity of multiple signaling pathways. However, because these pathways converge on common nodes, such as the induction of interferon, additional studies are needed to determine whether DUX4 inhibits unique components in each pathway or a common component responsible for ISG upregulation across pathways. We decided to focus further efforts on identifying the mechanism behind the suppression of IFNγ-mediated transcription, as this pathway was most broadly suppressed by DUX4.

### DUX4 transcriptional activity is not necessary for ISG suppression

There are two conserved regions of the DUX4 protein, the N-terminal homeodomains (aa19-78, aa94-153) and an ∼50 amino acid region at the end of the C-terminal domain (CTD) that is required for transcriptional activation by DUX4 (aa371-424) (Choi et al., 2016; Geng et al., 2011; Leidenroth & Hewitt, 2010). A mutation in the first homeodomain, F67A, prevents DUX4 DNA binding and target gene activation (Wallace et al., 2011). When expressed in MB135 cells, iDUX4-F67A did not activate the DUX4 target gene *ZSCAN4* yet still suppressed ISG induction by IFNγ (**Fig 2A and B**, and **Suppl Fig S2B**). A second construct, iDUX4aa154-424, which has the N-terminal homeodomain region replaced by a 3x FLAG tag and nuclear localization signals (3xFLAG-NLS) cassette (hereafter called iDUX4-CTD), was also transcriptionally silent yet equally suppressed activation of ISGs (**Fig 2A and C** and **Suppl Fig S2C**). RNA sequencing analysis using the same criteria to characterize ISG suppression by the full-length DUX4 demonstrated that the F67A mutant suppressed 70% of induced genes by more than 2-fold, or 41% of induced genes by more than 4-fold; whereas the iDUX4-CTD showed 90% or 52% suppression, respectively (**Suppl Table S1**). Together, these data indicate that DUX4 transcriptional activity is not necessary to suppress IFNγ-mediated gene induction.

**Figure 2.**
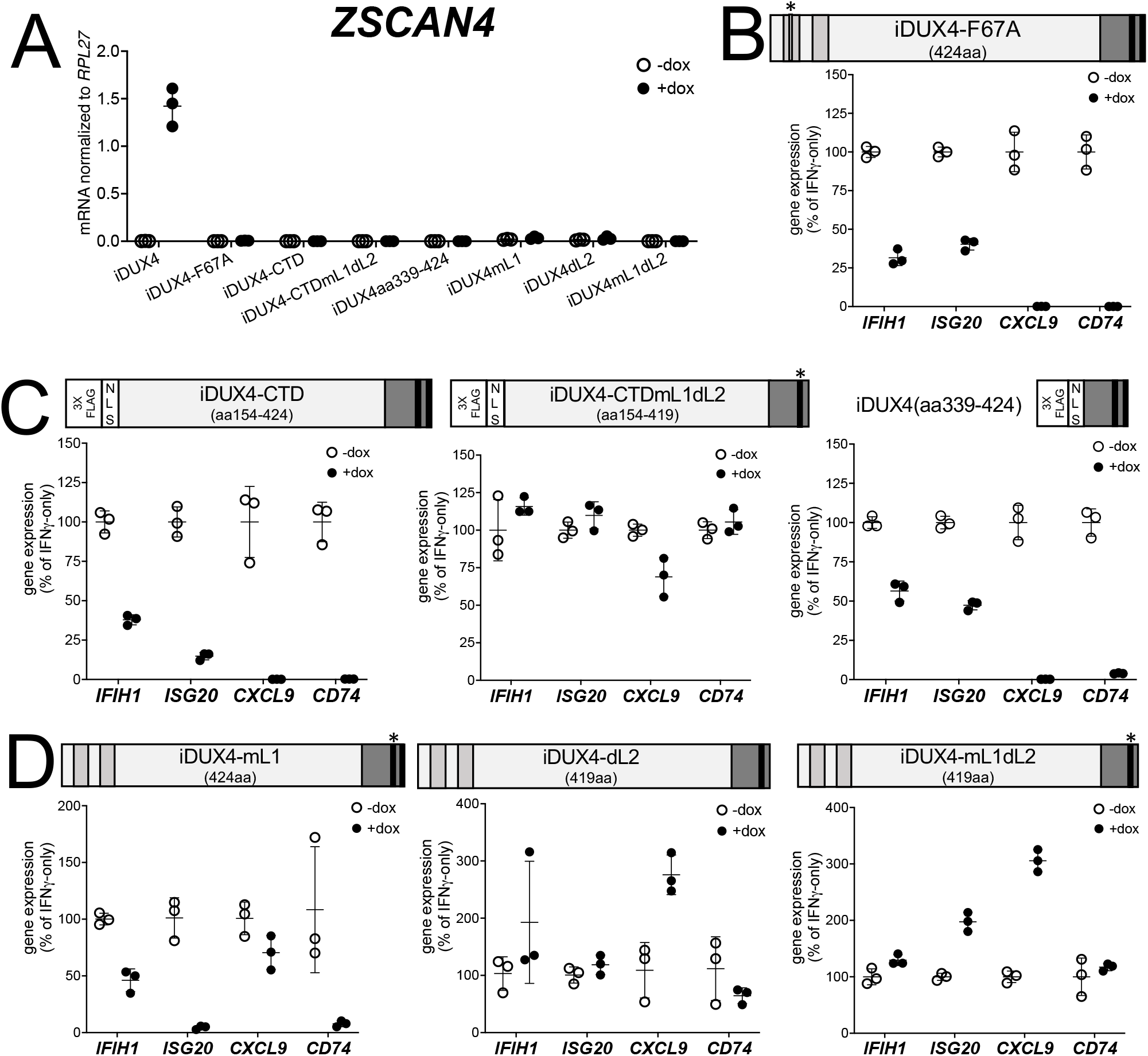
DUX4 transcriptional activity is not necessary for ISG suppression, whereas the C-terminal domain (CTD) is both necessary and sufficient. (**A**) MB135 cell lines with the indicated doxycycline inducible transgene ±doxycycline, were evaluated for *ZSCAN4* expression by RT-qPCR as a measure of the ability of the construct to activate a DUX4-target gene. Ct values were normalized to the housekeeping gene *RPL27*. Data represent the mean ±SD of three biological replicates with three technical replicates each. (**B-D**) MB135 cell lines with the indicated doxycycline inducible transgene were treated with IFNγ ±doxycycline. RT-qPCR was used to evaluate expression of *IFIH1, ISG20, CXCL9*, and *CD74* and Ct values were normalized to the housekeeping gene *RPL27*, then normalized to the IFNγ-only treatment to set the induced level to 100%. Data represent the mean ±SD of three biological replicates with three technical replicates each. Light gray, homeodomains; medium gray, conserved region of CTD; black, (L)LxxL(L) motifs; * indicates sites of mutation for F67A in HD1 and mutation of first LLDELL to AADEAA. **See Suppl Fig S2** for additional cell lines.

### The C-terminal Domain (CTD) is necessary and sufficient to suppress ISGs

The DUX4-CTD contains a pair of (L)LxxL(L) motifs, LLDELL and LLEEL, that are conserved across the DUXC/DUX4 family (Leidenroth & Hewitt, 2010). DUX4 transgenes with mutations in the first motif, deletion of the second motif, or both (iDUX4mL1, iDUX4dL2, iDUX4mL1dL2) (see **Suppl Fig 1B** for sequences of these mutants) failed to activate the DUX4 target *ZSCAN4* (**Fig 2A**). iDUX4ml1dl2 and iDUX4dl2 both lost the ability to suppress the panel of ISGs, whereas iDUX4mL1 showed partial activity, suppressing 3 of the 4 ISGs (**Fig 2D and Suppl Fig S2D**), indicating that these (L)LxxL(L) motifs are necessary for both ISG suppression and for transcriptional activation by DUX4.

To test sufficiency, we generated two additional C-terminal fragments of DUX4 (**Fig 2C**). The first, iDUX4-CTDmL1dL2, contains the CTD of iDUX4mL1dL2 with its N-terminal HDs replaced with the 3xFLAG-NLS cassette. Similar to iDUX4mL1dL2, iDUX4-CTDmL1dL2 did not block the panel of ISGs (**Fig 2C and Suppl Fig S2E)**. The second construct, iDUX4aa339-424, contains only the C-terminal 85 aa residues including both (L)LxxL(L) motifs, and maintained ISG suppression, though not as strongly on the *IFIH1* and *ISG20* genes (**Fig 2C and Suppl Fig S2F**). In summary, these data support a model in which the DUX4-CTD is both necessary and sufficient to suppress a major portion of the ISG response to IFNγ.

### The DUX4 protein interacts with STAT1 and additional immune response regulators

As an unbiased method to identify proteins that interact with the C-terminal region of DUX4, we conducted two experiments using liquid chromatography mass spectroscopy (LC-MS) to identify proteins that co-immunoprecipitated with DUX4-CTD constructs expressed in MB135 myoblasts. In the first experiment, we used MB135iDUX4-CTD cells either untreated, treated with doxycycline alone, or with both doxycycline and IFNγ. In the second experiment, we used MB135iDUX4-CTD and MB135iDUX4mL1dL2 cells both treated with doxycycline and IFNγ, compared to these two cell lines untreated and combined as a control. Proteins with a minimum of 2 peptide spectrum matches (PSMs) in at least one sample that were identified in both experiments were assigned to one of ten categories (see Methods) to separate candidate interactors from other categories that might be co-purified because of obligate interactions (e.g., proteasome or ribosome) or might be less likely to be relevant to immune responses (e.g., cytoskeletal proteins). Candidate interactors were then ranked based on the total PSMs for that protein across all samples. (It is important to note that the “bait” constructs were expressed at low levels in the samples not treated with doxycycline and that the immunoprecipitation concentrated this background, which might account for some of the candidate proteins appearing in the untreated samples.) STAT1 and DDX3X, two key regulators of innate immune signaling, ranked at the top of the list of candidate DUX4-CTD interactors, together with several other proteins implicated in modulating innate immune signaling (**Fig 3, left panel and Suppl Table S2**). Western blot analysis using independent biological samples from a co-IP experiment with MB135-iDUX4-CTD and MB135-iDUXB (as a control) validated the DUX4-CTD interactions with DDX3X, STAT1, PRKDC, YBX1, HNRNPM, PABPC1, NCL, CDK4, and HNRPU (**Fig 3, right panel**).

**Figure 3.**
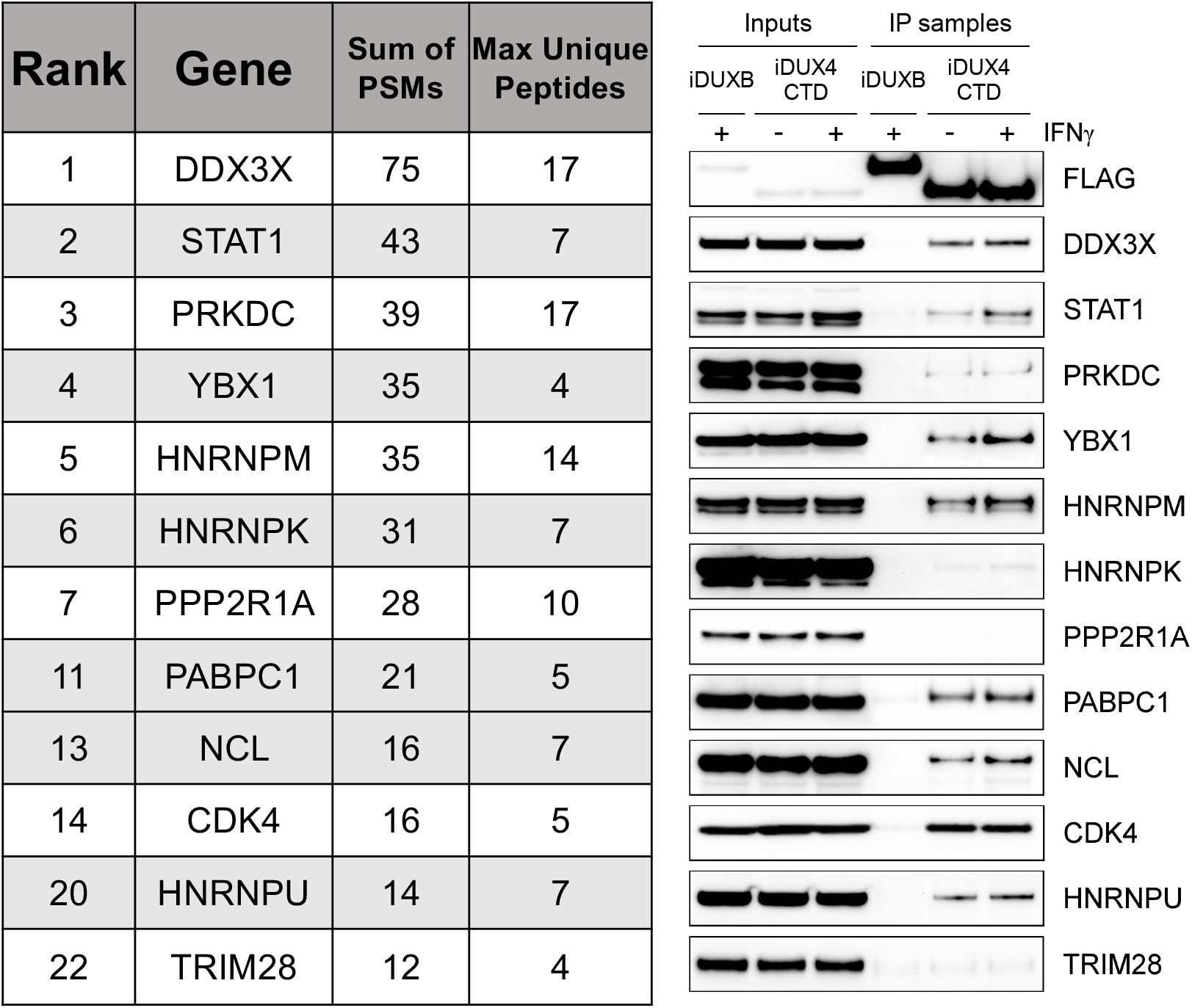
The DUX4 protein interacts with STAT1 and additional immune response regulators. Left panel, representative candidate interactors identified by mass spectrometry of proteins that co-immunoprecipitated with the DUX4-CTD and their relative ranking in the candidate list (see **Suppl Table S2** for full list). Right panel, validation western blot of proteins that co-immunoprecipitate with the DUX4-CTD in cell lysates from MB135 cells expressing doxycycline-inducible 3xFLAG-DUXB or 3xFLAG-DUX4-CTD, ±IFNγ treatment. Data represent biological duplicates. See Figure 3 Source Data for uncropped/raw images.

### The DUX4-CTD preferentially interacts with STAT1 phosphorylated at Y701

Because of its central role in IFNγ signaling, we elected to focus on the interaction of STAT1 with DUX4. To map the region(s) of the DUX4-CTD necessary to interact with STAT1, we expressed a truncation series in MB135 cells (all with an N-terminal 3xFLAG tag and NLS and all treated with IFNγ): iDUX4-CTD (aa154-424), iDUX4aa154-372, iDUX4aa154-308, and iDUX4aa154-271. The region of DUX4 between amino acids 271 and 372 was necessary for co-IP of STAT1, whereas the region between 372 and 424 containing the (L)LxxL(L) motifs might enhance DUX4-CTD binding to the phosphorylated forms of STAT1 (**Fig 4A**).

**Figure 4.**
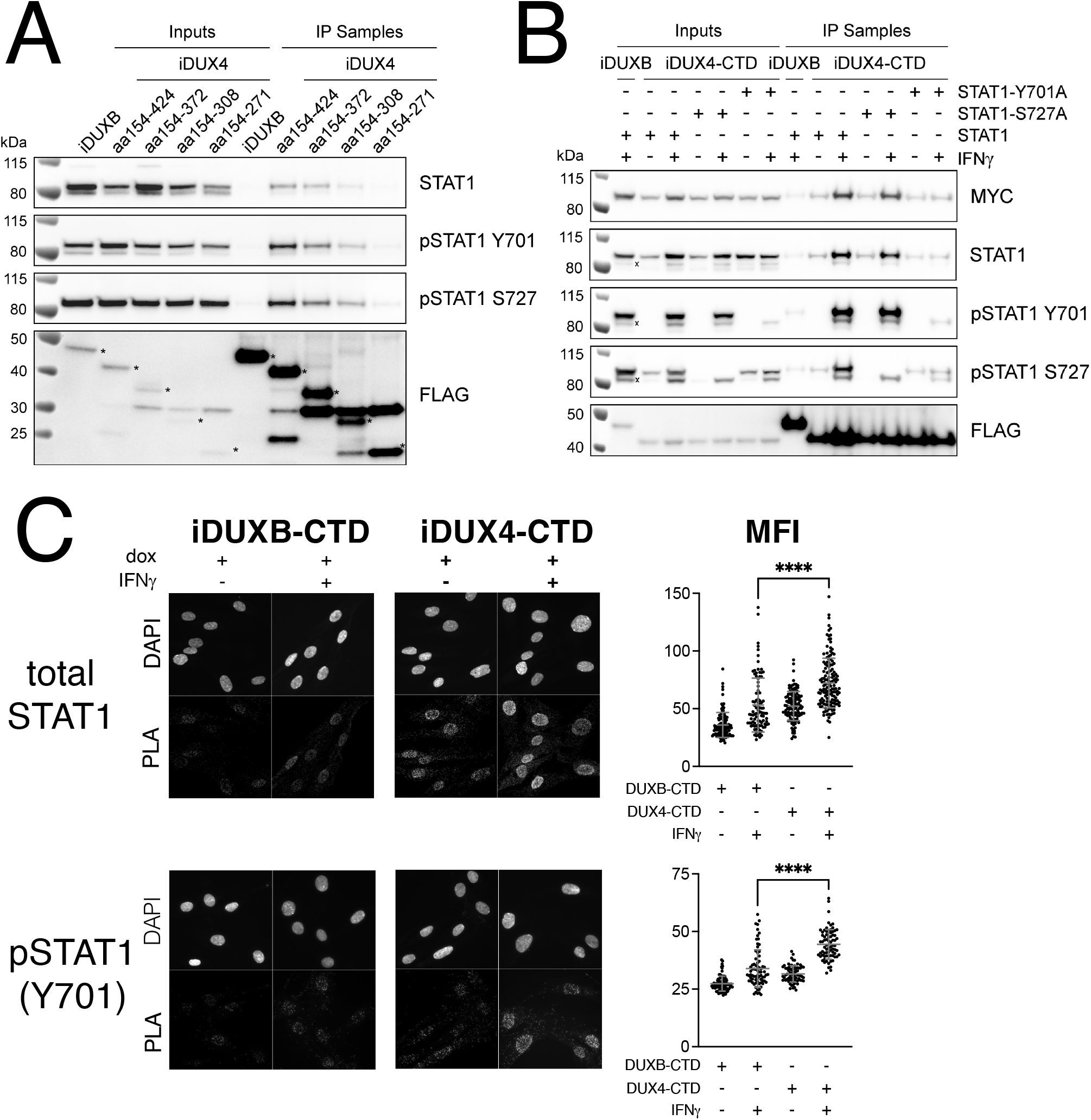
The DUX4-CTD preferentially interacts with pSTAT1-Y701. (**A**) Western blot showing input and immunoprecipitated proteins from either 3xFLAG-iDUXB (DUXB) or a truncation series of the 3x-FLAG-iDUX4-CTD cells (iDUX4) precipitated with anti-FLAG and probed with the indicated antibodies. Serial deletions of the iDUX4-CTD were assayed as indicated. All samples were treated with IFNγ. An asterisk indicates the correct band for each FLAG-tagged construct. See Figure 4a Source Data for uncropped/raw Western blots. (**B**) Input and anti-FLAG immunoprecipitation from 3xFLAG-iDUXB or 3x-FLAG-iDUX4-CTD cells co-expressing doxycycline inducible 3xMYC-iSTAT1, -iSTAT1-Y701A, or -iSTAT1-S727A with or without IFNγ treatment and probed with the indicated antibodies. An “x” indicates the endogenous (non-MYC tagged) STAT1 band. See Figure 4b Source Data for uncropped/raw Western blots. (**C**) Proximity-ligation assay (PLA) showing co-localization of endogenous STAT1 and pSTAT1 701 with the iDUX4-CTD compared to the interaction with the DUXB-CTD, in the nuclear compartment of IFNγ- and doxycycline-treated MB135 cells. Mean fluorescent intensity (MFI) of the nuclei in the PLA channel was measured for 10 images per cell line and treatment and plotted (**** p<0.0001, unpaired t-test). An unpaired t-test was used because the samples are biologically independent.

To determine whether phosphorylation of STAT1 enhanced interaction with DUX4, we co-expressed the FLAG-tagged iDUX4-CTD with a MYC-tagged iSTAT1 or STAT1 mutants Y701A or S727A, wherein doxycycline would induce expression of both the DUX4 and STAT1 transgenes. The wild-type STAT1 and STAT1-S727A showed enhanced binding to the CTD with IFNγ treatment, whereas IFNγ did not enhance the binding of STAT1-Y701A (**Fig 4B**). Furthermore, Immunofluorescence showed that DUX4-CTD expression did not prevent translocation of STAT1 to the nucleus following IFNγ treatment (**Suppl Fig S3**) and proximity Ligation Assay (PLA) indicated close interaction between the iDUX4-CTD and endogenous pSTAT1-Y701 in the nucleus of MB135 cells treated with doxycycline and IFNγ (**Fig 4C**). Therefore, the interaction between DUX4-CTD and STAT1 is enhanced by phosphorylation of STAT1-Y701 and this interaction happens within the nuclei of DUX4-CTD expressing cells.

### The DUX4-CTD decreases STAT1 occupancy at ISG promoters and blocks Pol-II recruitment

Chromatin immunoprecipitation (ChIP) was performed on MB135-iDUX4-CTD cells to assess STAT1 binding to ISG promoters. Compared to a gene-desert region where there should not be STAT1 binding (h16q21), there was a robust induction of STAT1 binding following IFNγ treatment at the promoters of several ISGs (*GBP1, IDO1, CXCL10*) with previously characterized STAT1 binding sites (Rosowski et al., 2014) (**Fig 5A, left four panels**). Treatment with IFNγ following induction of DUX4-CTD diminished STAT1 occupancy at all three ISGs, and paired RT-qPCR confirmed that the DUX4-CTD robustly suppressed the RNA induction by IFNγ (**Fig 5A, right panel)**. We used CUT&Tag (Cleavage Under Target & Tagmentation) (Kaya-Okur et al., 2019) to assess Pol-II occupancy genome wide and found that DUX4-CTD blocked Pol-II recruitment to ISGs without affecting occupancy at other genes (**Fig 5B**).

**Figure 5.**
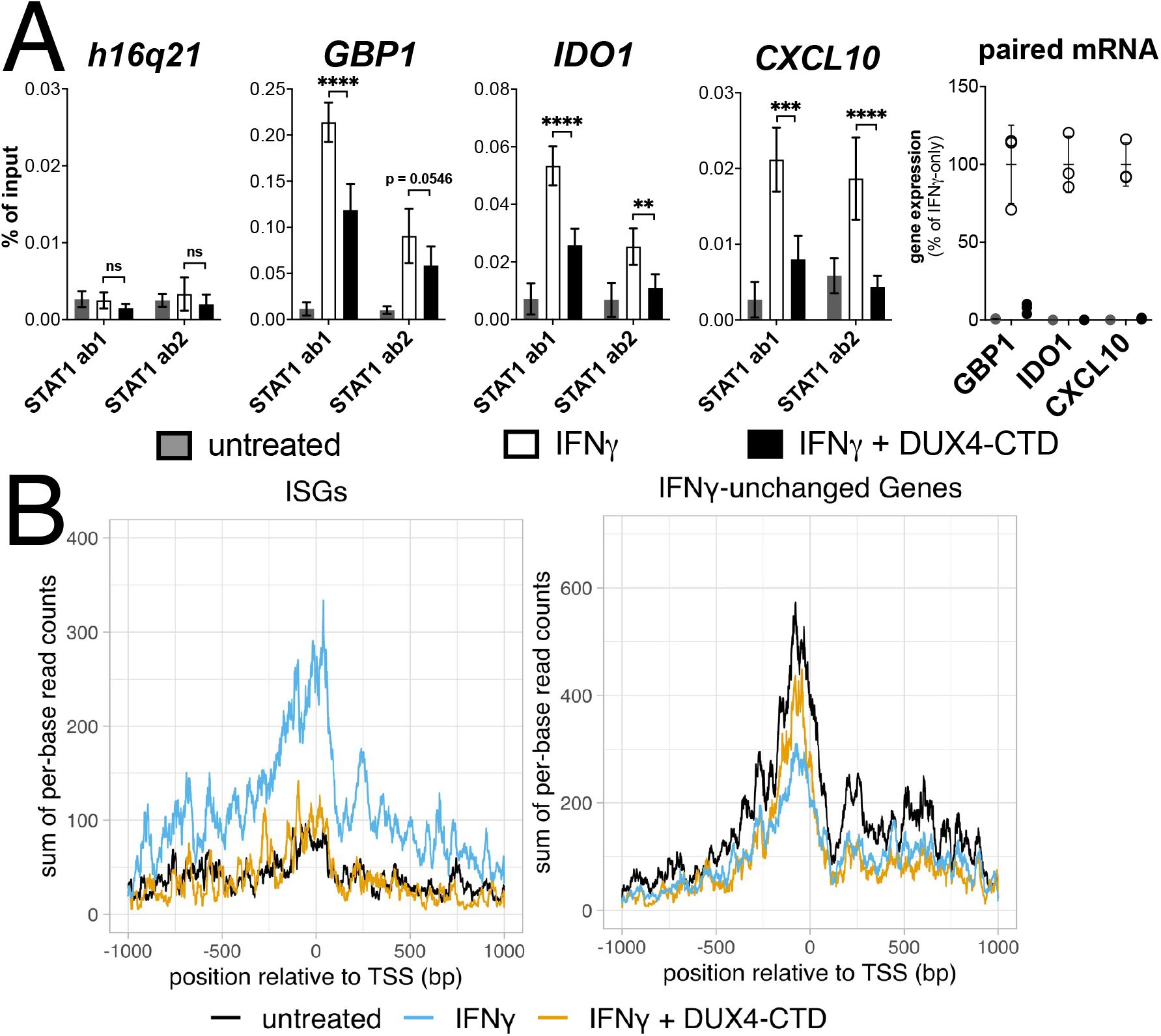
The DUX4-CTD decreases STAT1 occupancy at ISG promoters and blocks Pol-II recruitment. (**A, left four panels**) Chromatin immunoprecipitation using anti-STAT1 or IgG from MB135-iDUX4-CTD cells untreated, IFNγ-treated, or IFNγ and doxycycline treated. Ab1: 50:50 mix of STAT1 antibodies Abcam ab239360 and ab234400; Ab2: Abcam ab109320. ChIP-qPCR analysis relative to a standard curve constructed from purified input DNA was used to determine the quantity of DNA per IP sample, which was then graphed as a percent (%) of input. Data represent the mean ±SD of two biological replicates with 3 technical replicates each (**** p<0.0001, *** p<0.01, ** p<0.05, unpaired t-test). An unpaired t-test was used because the samples are biologically independent. (**A, right panel**) RT-qPCR of RNA from cells used for STAT1 ChIP showing induction of ISGs by IFNγ and suppression by DUX4-CTD. (**B**) CUT&Tag data showing the intensity of Pol-II signal across a 2000bp window centered on the TSS of ISGs (left) or IFNγ-unchanged genes (right) in untreated, IFNγ-treated, or IFNγ and doxycycline treated MB135-iDUX4-CTD cells.

### Endogenous DUX4 expression in FSHD myotubes is associated with suppressed ISGs

DUX4 expression in cultured FSHD muscle cells is often described as low, however this is due to high heterogeneity caused by strong expression in a small population of cells (Rickard et al., 2015; Snider et al., 2010). In cultured FSHD myotubes, approximately 5% of the myotubes might express DUX4 in their nuclei. To determine whether endogenous DUX4 suppresses IFNγ signaling, we assessed IFNγ induction of IDO1 in FSHD myotubes. Differentiation of FSHD myoblasts into multinucleated myotubes results in distinct populations of DUX4-expressing and DUX4-negative myotubes in the same culture, allowing for side-by-side evaluation of DUX4-positive and DUX4-negative muscle cells in the same culture. We determined the IFNγ induction of IDO1 as a representative ISG based on its low basal expression in skeletal muscle and our prior demonstration that it is suppressed in the MB135-iDUX4-CTD cells (see **Fig 5B, right panel**). Treatment with IFNγ produced a reliable IDO1 signal within the nucleus and cytoplasm of individual myotubes that did not express DUX4, whereas DUX4-positive myotubes did not show IDO1 expression in response to IFNγ (**Fig 6A**). Therefore, similar to our MB135-iDUX4 studies, endogenous DUX4 expressed at a physiological level is sufficient to prevent ISG induction by IFNγ.

**Figure 6.**
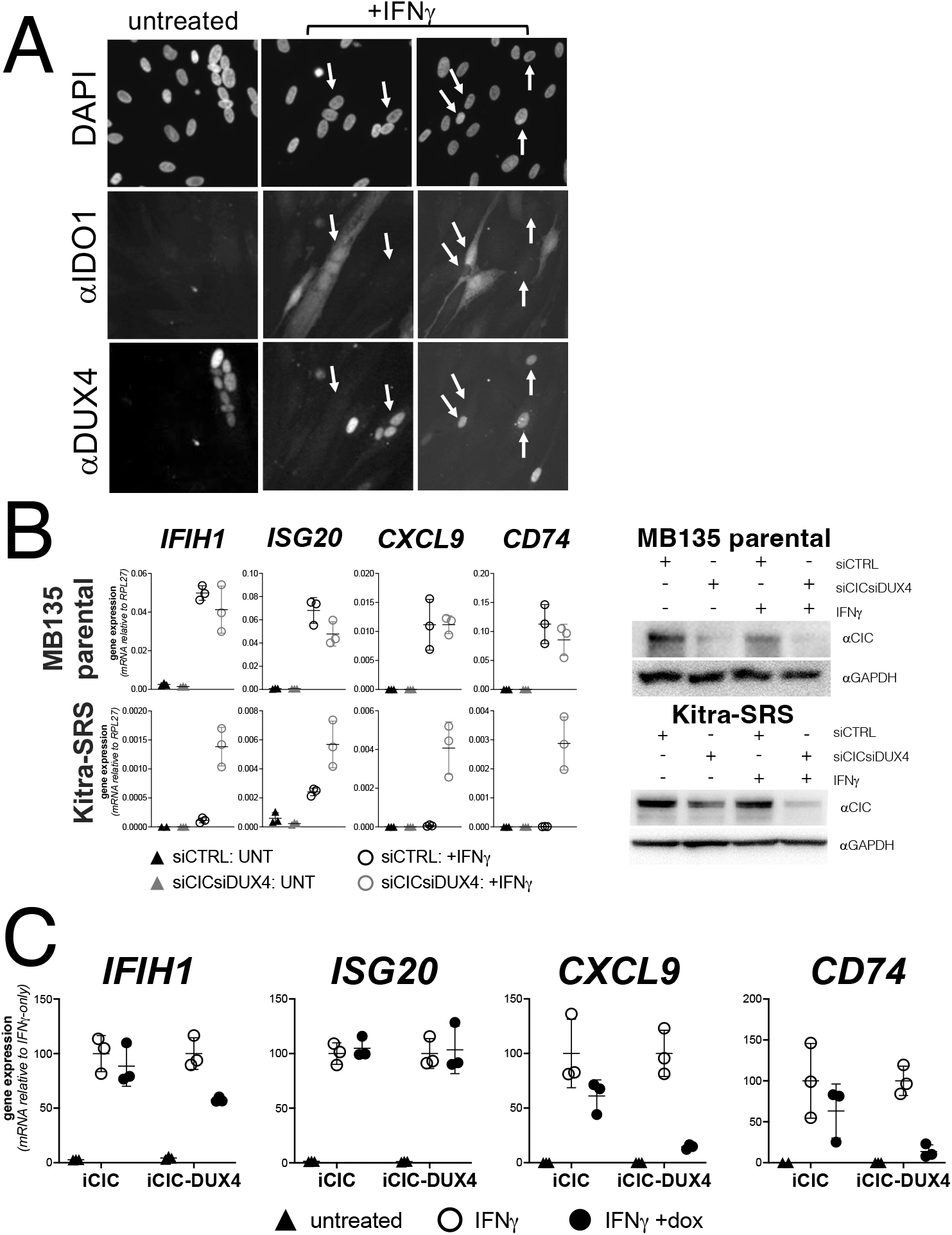
Endogenous DUX4 suppresses ISG induction in FSHD muscle cells and in a sarcoma cell line expressing a CIC-DUX4 fusion gene. (**A**) FSHD MB200 myoblasts were differentiated into myotubes, which results in the expression of endogenous DUX4 in a subset of myotubes. Cultures were treated ±IFNγ, and DUX4 and IDO1 were visualized by immunofluorescence. A representative image of DUX4+ and DUX4-myotubes shows IDO1 induction only in the DUX4-myotubes (arrows). (**B, left panel**) RT-qPCR of the indicated genes in MB135 parental or Kitra-SRS that express a CIC DUX4-fusion gene containing the DUX4 CTD. Cells were transfected with control or CIC- and DUX4-targeting siRNAs. Ct values were normalized to the house-keeping gene *RPL27*. Data represent the mean ±SD of three biological replicates with three technical replicates each. (**B, right panel**) Western blot showing lysates from MB135 or Kitra-SRS cells treated with control or CIC- and DUX4-targeting siRNAs ±IFNγ and probed with the indicated antibodies. See Figure 6b Source Data for uncropped/raw Western blots. (**C**) RT-qPCR of the indicated genes in MB135 with an inducible CIC (MB135-iCIC) or an inducible CIC-DUX4 fusion gene (MB135-iCIC-DUX4). Cells were untreated, IFNγ-treated, or IFNγ and doxycycline treated. Ct values were normalized to the housekeeping gene *RPL27*, then normalized to the IFNγ-only treatment to set the induced level to 100%. Data represent the mean ±SD of three biological replicates with three technical replicates each.

### Endogenous CIC-DUX4 fusion gene suppresses ISG induction in a sarcoma cell line

The majority of EWSR1 fusion-negative small blue round cell sarcomas have a genetic re-arrangement between CIC and DUX4 that creates a fusion protein containing the carboxyterminal (L)LxxL(L) motif region of DUX4 (Graham et al., 2012; Kawamura-Saito et al., 2006). We confirmed that the Kitra-SRS sarcoma cell line expresses a CIC-DUX4 fusion mRNA containing the terminal 98 amino acids of DUX4 as previously described (Nakai et al., 2019). Compared to MB135 myoblasts, Kitra-SRS cells showed absent-to-low induction of ISGs when treated with IFNγ and control siRNAs. In contrast, siRNA knockdown of the CIC-DUX4 fusion in the KitraSRS cells resulted in a substantially increased IFNγ-induction of ISGs, whereas knockdown of CIC in the MB135 cells did not alter ISG induction **(Fig 6B)**. To confirm that the CIC-DUX4 fusion was suppressing ISG induction, we expressed a doxycycline inducible CIC or the Kitra-SRS CIC-DUX4 fusion protein in MB135 cells and showed that the CIC-DUX4 fusion, but not CIC, suppressed IFNγ-induction of ISGs *IFIH1, CXCL9*, and *CD74*, although not *ISG20* (**Fig 6C**).

### Conservation of ISG repression and STAT1 interaction in mouse Dux

*Dux*, the mouse ortholog of human *DUX4*, is expressed at the equivalent developmental stage to human *DUX4* (Hendrickson et al., 2017), activates a parallel transcriptional program (Whiddon et al., 2017), and contains the (L)LxxL(L) motifs that we have shown to be necessary for ISG repression by human DUX4. In fact, the mouse Dux sequence contains a 60 amino acid triplication of the (L)LxxL(L)-containing region (**Supp Fig S4**). Accordingly, we introduced a doxycycline-inducible mouse *Dux* transgene into human MB135 cells (MB135-iDux) and found that the full-length Dux protein repressed the panel of ISGs even more robustly than the full-length or CTD portion of human DUX4 (**Fig 7A**). Similar to human DUX4, Western analysis confirmed the co-immunoprecipitation of STAT1 and both phosphorylated pSTAT1-Y701 and pSTAT1-S727 with mouse Dux (**Fig 7A**). These data demonstrate that the suppression of ISG induction and interaction with phosphorylated STAT1 is conserved in the DUXC family.

**Figure 7.**
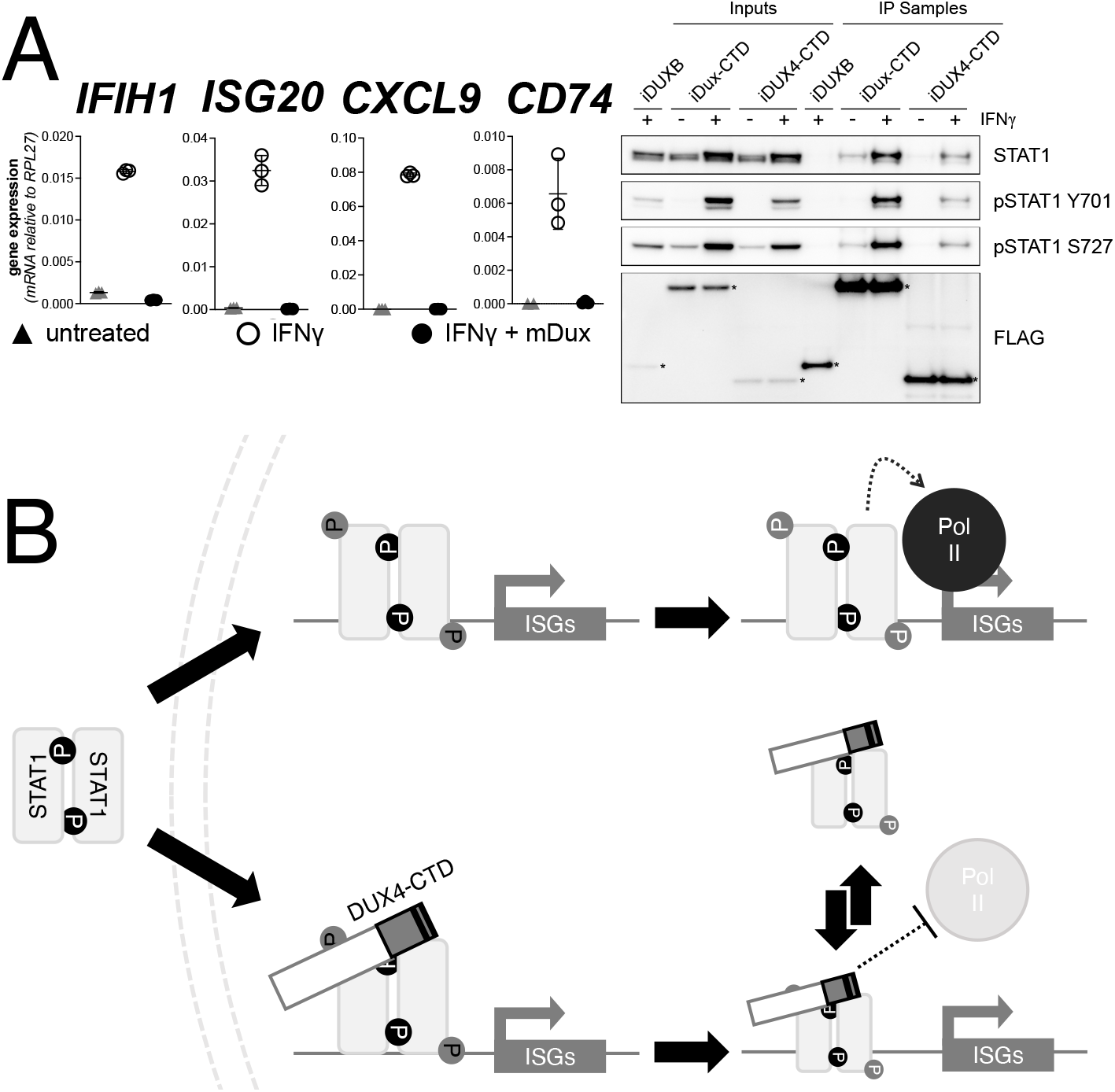
Conservation of ISG repression and STAT1 interaction in mouse Dux. (**A, left panel**) RT-PCR of the indicated genes in MB135-iDux cells untreated or treated with IFNγ ±doxycycline. Ct values were normalized to the housekeeping gene *RPL27*, then normalized to the IFNγ-only treatment to set the induced level to 100%. Data represent the mean ±SD of three biological replicates with three technical replicates each. (**A, right panel**) Western blot showing input and immunoprecipitated proteins from either 3xFLAG-iDux or 3x-FLAG-iDUXB cells ±IFNγ precipitated with anti-FLAG and probed with the indicated antibodies. See Figure 7a Source Data for uncropped/raw Western blots. (**B**) A model supported by the data showing how the DUX4-CTD might prevent STAT1 ISG induction. (Top) In the absence of the DUX4-CTD, pSTAT1 Y701 (black “P”) dimerizes, translocates to the nucleus, binds its GAS motif in the ISG promoter, acquires secondary phosphorylation at S727 (grey “P”), and recruits a stable transcription complex that includes Pol II to drive transcription of ISGs. (Bottom) In the presence of the DUX4-CTD, STAT1 is phosphorylated, translocates to the nucleus, and binds its GAS motif as evidenced by the pSTAT1 S727 in complex with the CTD. However, diminished steady-state occupancy of STAT1 at the ISG promoters and absence of Pol-II recruitment indicate that the STAT1-DUX4-CTD complex does not stably bind DNA and fails to recruit Pol-II and the pre-initiation complex. The (L)LXXL(L) motifs (black bars in DUX4-CTD) are necessary to interfere with transcription suppression and likely prevent STAT1 from interacting with a factor in the pre-initiation complex or recruit a co-repressor.

## DISCUSSION

In this study, we show that the DUX4-CTD, a transcriptionally inactive carboxyterminal fragment of DUX4, is necessary and sufficient to broadly suppress ISG induction by IFNγ as well as partially inhibit induction through the IFNβ, cGAS, IFIH1/MDA5, and DDX58/RIG-I pathways. The DUX4-CTD colocalizes with STAT1 in the nucleus, diminishes steady-state STAT1 occupancy at ISG promoters, and prevents Pol-II recruitment and transcriptional activation of ISGs by IFNγ. Whereas the conserved DUX4 (L)LxxL(L) motifs are necessary to suppress transcriptional activation by STAT1, they are not necessary for the interaction of DUX4 and STAT1. The suppression of IFNγ signaling by endogenous DUX4 in FSHD muscle cells and the CIC-DUX4 fusion protein in sarcomas provides support for the biological relevance of these findings.

Our data support a simple model of how DUX4 inhibits STAT1 activity (**Fig 7B**). IFNγ binding to its receptor, IFNGR, leads to the phosphorylation of STAT1 at Y701, subsequently STAT1 forms a homodimer, translocates to the nucleus, and binds the gamma-activated sequence (GAS) in the promoters of ISGs. DNA-bound STAT1 is additionally phosphorylated at S727 and recruits Pol-II to the ISG promoters (Sadzak et al., 2008; Wen et al., 1995). Our studies show that DUX4-CTD interacts with STAT1 phospho-Y701 in the absence of phospho-S727 (i.e., binds the S727A STAT1 mutant), yet also efficiently co-immunoprecipitates with STAT1 phospho-S727 from cell lysates. This indicates that despite DUX4 interacting with STAT1 phospho-Y701, DNA binding of this complex is not fully impaired because of the association with STAT1 phospho-727. However, our ChIP and CUT&Tag studies show decreased STAT1 steady-state occupancy of ISG promoters and failure to recruit Pol-II. Together, these data support a model of DUX4 interaction with pSTAT1-Y701 that prevents the formation of a stable DNA-bound complex and recruitment of Pol-II, but likely not the initial binding of STAT1 to DNA because of the abundance of phospho-S727 associated with DUX4. The (L)LxxL(L) motifs are necessary to prevent transcriptional activation, presumably by blocking Pol-II recruitment, but not necessary for the interaction of DUX4 with STAT1. This could be due to recruitment of a repressor, or by simply blocking the interaction of STAT1 with an intermediate factor necessary to recruit Pol-II.

The (L)LxxL(L)-dependent inhibition of STAT1 by DUX4 in the current study bears a striking similarity to the inhibitory mechanisms displayed by LxxLL-containing members of the PIAS family. LxxLL motifs were first identified in nuclear-receptor (NR) signaling pathways (Heery et al., 1997) where they were found to facilitate protein-protein interactions between unbound NRs and co-repressors such as RIP140 and HDACs, or agonist-bound NRs and co-activators such as CBP/p300 (Plevin et al., 2005; Savkur & Burris, 2004). LxxLL motifs have since been characterized in multiple protein families, including the PIAS family, and specifically implicated in modulating immune transcriptional networks via interaction with and inhibition of STATs, IRFs, and NF-kB (Shuai & Liu, 2005). While the (L)LxxL(L) region of DUX4 is required for suppression of IFNγ-mediated ISG induction and its enhanced interaction with pSTAT1-Y701, it is not required for its apparently weaker interaction with unphosphorylated STAT1. In a similar manner, the LxxLL motif of PIASγ is not required for initial binding to STAT1, but is required to suppress ISG induction mediated by STAT1 in response to both IFNβ (Kubota et al., 2011) and IFNγ (Liu et al., 2001). The same motif is required for the trans-repression of androgen receptor (AR) signaling by PIASγ (Gross et al., 2001) and of Erythroid Krüppel-like factor (EKLF or KLF1) by PIAS3 (Siatecka et al., 2015), though again it is not required for the initial interaction of either pair. The studies referenced above hypothesize that this trans-repression relies on the recruitment of co-repressors, although the specific interactors were not determined. Additionally, just as DUX4 reduces the steady-state occupancy of STAT1 to DNA, PIAS proteins can suppress transcriptional networks by blocking DNA binding, as with PIAS3 and STAT3 (Chung et al., 1997) or PIAS1 and NF-kB p65 (Liu et al., 2005). These studies describe mechanisms of transcriptional suppression by LxxLL motifs in PIAS and other proteins that have strong parallels to the (L)LxxL(L) motifs in human DUX4 and mouse Dux. It is important to emphasize that the xx amino acids in the DUXC family are acidic and there is conservation of flanking amino acids as well, suggesting that the DUXC family likely evolved target specificity through these larger areas of conservation.

In addition to STAT1, the mass spectrometry identified several proteins that interact with the DUX4-CTD that might also have a role in modulating immune signaling. Although additional work is needed to validate the biological relevance of these interactions, many have functions related to immune signaling and that will need to be evaluated in future studies. DDX3X and PRKDC are the top ranked candidates, together with STAT1. DDX3X has been shown to regulate RNA processing, translation, and innate immune signaling (Mo et al., 2021). It was also shown to be a pathway specific regulator of IRF3 and IRF7 in part by acting as a scaffolding factor necessary for IKK-ε and TBK1 phosphorylation of IRFs (Gu et al., 2013; Schroder et al., 2008). DDX3X was also shown to be a sensor of dsRNA and viral stem-loop RNA with a role in the initial induction of ISGs, including IFIH1 and DDX58 (Oshiumi et al., 2010) that then serve to amplify the signaling mechanisms. PRKDC is known mostly for its major roles in DNA repair but also has been implicated in regulating the response to cytoplasmic DNA through the cGAS and IRF3 pathway (Ferguson et al., 2012).

Our current findings also provide a molecular mechanism for the suppression of IFNγ stimulated genes in DUX4-expressing cancers. Previously we reported that the full-length DUX4 is expressed in a diverse set of solid cancers (Chew et al., 2019). Cancers expressing DUX4 had diminished IFNγ-induced MHC Class I expression, reduced anti-tumor immune cell infiltration, and showed resistance to immune checkpoint blockade. In our current study, we show that the CIC-DUX4 fusion in EWSR1-fusion-negative sarcomas blocks IFNγ-induced ISG expression. This fusion protein contains the terminal 98 amino acids of DUX4, aa327-424, that encompasses a region shown to be sufficient to suppress IFNγ signaling in the iDUX4-aa339-424 (see **Fig 2C**). It is reasonable to suggest that this fusion protein in the CIC-DUX4 sarcomas, or the full length DUX4 in some other cancers, contributes to immune evasion at least in part through its interaction with STAT1, and that targeting DUX4 or its interaction with STAT1 might improve immune-based therapies for DUX4-expressing cancers.

The conservation of the (L)LxxL(L) motifs in mouse Dux and its similar interaction with STAT1 and inhibition of IFNγ signaling indicates that this is a conserved function of the DUXC family. DUX4, Dux, and the canine DUXC all induce expression of endogenous retroelements, as well as pericentromeric satellite repeats that form dsRNAs that, at least in the case of DUX4, induce a dsRNA response that results in activation of PKR and phosphorylation of EIF2α (Shadle et al., 2019; Shadle et al., 2017). Therefore, it is possible that the interaction with STAT1 and other immune signaling modulators might prevent the activation of the ISG pathway while permitting the PKR response, although the biological consequences remain to be further explored. It is also interesting that DUX4, Dux and possibly other members of the DUXC family are expressed in immune privileged tissues—i.e., cleavage embryo, testis and thymus—and our study suggests that their expression might contribute to this immune privileged state.

## MATERIALS AND METHODS

### Cell culture

All myoblast experiments were conducted in immortalized MB135 (*H. sapiens*, Female, control, Fields Center for FSHD and Neuromuscular Research at the University of Rochester Medical Center, https://www.urmc.rochester.edu/neurology/fshd-center.aspx) or MB200 (*H. sapiens*, Male, FSHD2 subject, Fields Center for FSHD and Neuromuscular Research at the University of Rochester Medical Center, https://www.urmc.rochester.edu/neurology/fshd-center.aspx) cell lines, respectively cultured in Ham’s F-10 Nutrient Mix (Gibco) supplemented with 15% fetal bovine serum (Hyclone), 100 U/100 µg/ml penicillin/streptomycin (Gibco), 1µM dexamethasone (Sigma), and 10ng/mL recombinant human basic fibroblast growth factor (PeproTech). To differentiate the myoblasts to myotubes, media was changed to DMEM supplemented with 10 ug/ml insulin (Sigma) and 10 ug/ml transferrin (Sigma). Cell lines containing doxycycline-inducible transgenes were additionally cultured with 2 µg/mL puromycin (Sigma). Transgenes were induced with 1 µg/mL of doxycycline (Sigma) for 4 hours prior to other treatments for a total of 20 hrs. The Kitra-SRS cells (RRID:CVCL_YI69) were provided by Dr. H. Otani and Osaka University (Nakai et al., 2019) and were cultured in DMEM supplemented with 10% fetal bovine serum (Hyclone) and 100 U/100 µg/ml penicillin/streptomycin (Gibco). Biological replicates consisted of independent but parallel experiments, such as simultaneously stimulating three cell culture plates with IFNγ. Technical replicates consisted of repeat measurements of the same biological sample, such as loading the same biological sample in triplicate for analysis by RT-qPCR.

### Cloning, virus production, and monoclonal cell line isolation

Human DUX4 and mouse Dux truncation constructs were created by cloning synthesized, codon-optimized gBlock fragments into the pCW57.1 vector (Addgene plasmid #41393) downstream of the doxycycline-inducible promoter. Lentiviral particles were created by transfecting 293T cells with the subcloned pCW57.1 expression vectors, psPAX2 (Addgene plasmid #12260), and pMD2.G (Addgene plasmid #12259) using Lipofectamine 2000 according to the manufacturer’s instructions (Invitrogen). Myoblasts were transduced and selected using 2 µg/mL puromycin at low enough confluence to allow for isolation of clonal lines using cloning cylinders. Transgenic clonal lines were validated for protein size, expression level, and localization by western blot and immunofluorescence.

### Immune stimulation and RT-qPCR

Myoblasts were transfected with either (final concentrations) 10 µM 2’,3’-cGAMP (Sigma Aldrich), 2 µg/mL poly(I:C) (Invitrogen), or 1 µg/mL 3’ppp-dsRNA RIG-I ligand (Dan Stetson Lab, UW) using Lipofectamine 2000 according to manufacturer’s protocol, or were stimulated with 1000U IFNβ (PeproTech) or 200 ng/mL IFNγ (R+D Systems) by addition directly to cell culture medium. After 16 hours of immune stimulation, RNA was collected from cells using the NucleoSpin RNA Kit (Macherey-Nagel) according to manufacturer’s instructions. RNA samples were quantified by nanodrop and 1 µg of RNA per sample was treated with DNase I Amplification Grade (Thermo Fisher), and then synthesized into cDNA using the Superscript IV First-Strand Synthesis System, including oligo dT primers (Invitrogen). qPCR was run in 384-well plates on an Applied Biosystems QuantStudio 6 Flex Real-Time PCR System (ABI) and analyzed in Microsoft Excel.

### RNA-seq Library Preparation and Sequencing

RNA was extracted as described above from untreated, doxycycline-treated, IFNγ-treated, or doxycycline- and IFNγ-treated samples. RNA was submitted to the Fred Hutchinson Cancer Research Center Genomics Core for library preparation using the TruSeq3 Stranded mRNA kit (Illumina) followed by size and quality analysis by Tapestation (Agilent). Libraries were sequenced on a NextSeq P2-100 (Illumina).

### RNA-seq Analysis

Sequencing analysis was performed using R version 4.0.3 (R Core Team, 2020). Sequencing reads were trimmed using Trimmomatic (version 0.39) (Bolger et al., 2014), and aligned to the Homo sapiens GRCh38 reference genome with the Rsubread aligner (Liao et al., 2019). Gene counts were analyzed using featureCounts (v2.0.1) (Liao et al., 2019) and the Gencode v35 annotation file. Normalization and differential expression analysis were done with DESeq2 (v1.26.0) (Love et al., 2014).

### Immunofluorescence

Cells were fixed for 10 minutes with 2% paraformaldehyde (Thermo Scientific) for DUX4/STAT1 and 4% paraformaldehyde for DUX4/IDO1 then permeabilized for 10 minutes with 0.5% Triton X-100 (Sigma), both at room temperature with gentle shaking. Cells were then blocked for 2 hours with PBS/0.3M glycine/3% BSA at room temperature with gentle shaking. Primary antibodies were incubated at 4°C overnight: rabbit anti-IDO1 (D5J4E) 1:100 (Cell Signaling Tech 86630S, RRID: AB2636818), mouse anti-DUX4 (P2G4) 1:250 (Geng et al., 2011), mouse anti-FLAG M2 1:500 (Sigma #F1804, RRID: AB_262044), rabbit anti-STAT1 mAb 1:750 (Abcam #ab109320, RRID: AB_10863383). Cells were washed three times with 1X PBS containing 3% BSA, then secondary antibodies were incubated for 1hr at room temperature: FITC-conjugated donkey anti-rabbit (Jackson ImmunoResearch #711-095-152, RRID: AB_2315776) or TRITC-conjugated donkey anti-mouse (Jackson ImmunoResearch #715-025-020, RRID: AB_2340764). Cells were washed once with 1X PBS containing 3% BSA then stained with DAPI (Sigma) for 10’ at room temperature and visualized.

### Fractionated anti-FLAG immunoprecipitation

Cells were lysed on the plate with digitonin lysis buffer pH 7.4 (37.5 µg/mL digitonin, 25 mM Tris-HCl pH 7.5, 125 mM NaCl, 1 mM EDTA, 5% glycerol) supplemented with Pierce Protease Inhibitors EDTA-free (PIA32955) and Pierce Phosphatase Inhibitors (PIA32957), transferred to a centrifuge tube and incubated for 10 minutes at 4°C with rotation. Centrifugation at 2500 rcf at 4°C for 5 min pelleted the nuclei, supernatant was discarded, and nuclei resuspended in 1mL IP buffer pH 7.4 (25 mM Tris-HCl pH 7.5, 175 mM NaCl, 1 mM EDTA, 0.2% NP-40, 5% glycerol) and incubated for 1 hour at 4°C with rotation then spun at 21000 rcf for 10 minutes at 4°C to pellet insoluble debris. Protein concentration was determined using the Pierce BCA Protein Assay Kit (ThermoFisher, 23225). An equivalent amount of protein per sample was pre-cleared with Dynabeads Protein G (Invitrogen) bound to mouse IgG (Abcam #131368, RRID: AB_2895114) for 1 hour at 4°C with rotation. FLAG-tagged constructs were then immunoprecipitated with Dynabeads Protein G beads coupled to mouse anti-FLAG M2 mAb (Sigma #F3165, RRID:AB_259529) for 3 hours at 4°C with rotation. Beads were washed 3X with 1 mL IP buffer and eluted by adding 2X NuPage LDS Sample Buffer (Thermo Fisher, diluted from 4X with PBS) to the beads and heating for 10 minutes at 70°C.

### Liquid Chromatography Mass Spectroscopy (LC-MS)

For LC-MS, anti-FLAG immunoprecipitation was performed with beads cross-linked to the anti-FLAG antibody (Sigma #F3165, RRID:AB_259529) and the proteins competitively eluted with FLAG peptide. Eluted protein samples were electrophoresed into a NuPage 4-12% Bis-Tris gel, excised, and processed by the Fred Hutchinson Cancer Research Center Proteomics Core. Samples were reduced, alkylated, digested with trypsin, desalted, and run on the Orbitrap Eclipse Tribid Mass Spectrometer (Thermo Fisher). Proteomics data were analyzed using Proteome Discoverer 2.4 against a Uniprot human database that included common contaminants using Sequest HT and Percolator for scoring. Results were filtered to only include protein identifications from high confidence peptides with a 1% false discovery rate. Proteins that were identified in at least one sample from both independent experiments with at least 2 PSMs in one sample were assigned to one of ten categories: 1, candidates; 2, cytoskeletal associated; 3, cytoskeletal; 4, ribosome/translation associated; 5, proteasome associated; 6, membrane or extracellular; ER, golgi, or vesicle associated; 8, lipid metabolism; 9, chaperones; 10, nuclear import or nuclear membrane associated. The proteins in category 1 were further investigated for interactions with DUX4. It should be noted that this category assignment process de-prioritized groups of proteins based on assignment to a cellular compartment or function (e.g. ribosome/translation proteins might associate with DUX4 as part of a translation complex rather than having a role in immune signaling) and it is possible that some of the proteins assigned to the non-candidate categories might be functional interactors with DUX4 and have an important biological role.

### Chromatin immunoprecipitation and sequencing

Chromatin immunoprecipitation (ChIP) was performed as previously described (Nelson *et al* 2006) with the following modifications: Cells were plated and allowed to grow to 70-80% confluence. Cells were fixed with 1.42% formaldehyde for 15min at room temperature with shaking. Fixation was quenched with 125mM glycine, and cells were scraped into Falcon tubes and collected by centrifugation. Cells were lysed to isolate nuclei for 10min on ice using IP Buffer (150mM NaCl, 50mM Tris-HCl pH 7.4, 5mM EDTA, 1% Triton X-100, 0.5% NP-40) containing Pierce Protease Inhibitors EDTA-free (PIA32955) and Pierce Phosphatase Inhibitors (PIA32957) added fresh. Pelleted nuclei were sonicated on a Diagenode Biruptor on “Low” for 10 min as 30 sec on/30 sec off, followed by 4 rounds of sonication on “High” for 10 min each as 30 sec on/30 sec off (50 minutes total sonication) in IP Buffer + 0.5% SDS. For immunoprecipitation, 500 ng of chromatin was set aside per condition as an “Input” and 4 µg of antibody was added to 10 µg of chromatin in an equal volume of IP Buffer + 0.5% SDS across samples. “STAT1 Ab1” consisted of a 50:50 mix of rabbit anti-STAT1 [EPR21057-141] (Abcam #ab234400) and rabbit anti-STAT1 [EPR23049-111] (Abcam ab#239360). “STAT1 Ab2” was rabbit anti-STAT1 (Abcam #ab109320, RRID:AB_10863383). For an IgG control we used purified Rabbit Polyclonal Isotype Control Antibody (Biolegend #CTL-4112). IP Buffer was added to lower the percentage of SDS < 0.1%, and tubes were incubated with rotation overnight at 4°C. During this time, protein-A agarose Fastflow beads (Millipore) were washed twice with IP Buffer and then blocked in IP Buffer containing 2% BSA by rotating overnight at 4°C. After clearing the chromatin as described, beads were aliquoted to fresh tubes and the top 90% of chromatin was transferred to the tubes containing the blocked bead slurry. Tubes were rotated for 1 hour at 4°C. Beads were washed 5 times with cold IP Buffer containing 0.1% SDS, 2 times with cold IP Buffer containing 500mM NaCl, and 2 times with cold PBS. DNA was isolated as described in the original protocol and used as a template in qPCR. Input DNA was used to create a standard curve. qPCR primers for the h16q21 gene desert region and the ISGs were previously published (Maston et al., 2012; Rosowski et al., 2014).

### Proximity Ligation Assay

Cells were fixed for 10 minutes with 4% paraformaldehyde (Thermo Scientific), permeabilized for 10 minutes with 0.5% Triton X-100 (Sigma), and then blocked for 2 hours at room temperature with PBS/0.3M glycine/3% BSA. Primary antibodies were diluted in PBS/3% BSA and incubated with samples overnight at 4°C: anti-FLAG [M2] (F1804) (1:4000), anti-STAT1 [EPR4407] (1:1000), and anti-pSTAT1 Y701 [58D6] (1:1000). Samples were washed 3 times for 10 minutes with 1x Wash Buffer A (10mM Tris, 150mM NaCl, 0.05% Tween, adjusted pH to 7.4), and then incubated with Duolink In Situ PLA Probe Anti-Rabbit PLUS (Sigma, Cat# DUO92002) and Duolink In Situ PLA Probe Anti-Mouse MINUS (Sigma, Cat# DUO92004) diluted in PBS/3% BSA for 1 hour in a humidity chamber at 37°C. Samples were washed 3 times for 10 minutes with 1x Wash Buffer A, and then treated with ligase from the Duolink In Situ Detection Reagents Green kit (Sigma, Cat# DUO92014) for 30 minutes in a humidity chamber at 37°C. Samples were washed 3 times for 10 minutes with 1x Wash Buffer A, and then treated with polymerase from the Duolink In Situ Detection Reagents Green kit for 1 hour and 40 minutes in a humidity chamber at 37°C. Samples were washed 2 times for 10 minutes with 1x Wash Buffer B (200mM Tris, 100mM NaCl, adjusted pH to 7.5) and then once for 1 minute with 0.01x Wash Buffer B. Samples were mounted with Prolong Glass Antifade Mountant (ThermoFisher, Cat# P36983), and then visualized with a fluorescent microscope using FITC and DAPI filters.

### CUT&Tag

CUT&Tag was performed as previously described (Kaya-Okur et al., 2019) with the following modifications: MB135-iDUX4-CTD myoblasts were plated and allowed to grow to 70-80% confluence. Cells were left untreated, treated with 200ng/mL IFNg for 16hr, or pre-treated with 1µg/mL doxycycline for 4hr then had IFNg added directly to cell media for an additional 16hr. Fresh cells were harvested and washed in PBS, crosslinked with 0.1% formaldehyde for 90 seconds, then counted and 1.25e6 cells were aliquoted per reaction tube. *Drosophila* S2 cells were spiked-in at a genomic ratio of 1:10. Nuclei were prepared from cells in Buffer NE1 (20mM HEPES-KOH pH7.9, 10mM KCl, 0.1% Triton X-100, 20% glycerol, 0.5mM spermidine, Pierce Protease Inhibitors EDTA-free [PIA32955]) on ice for 10min and then bound to concanavalin A-coated beads for 10min. Primary antibody (dilution 1:50) was bound overnight at 4°C in 25µL per sample of Antibody Buffer (20mM HEPES-KOH pH7.5, 150mM NaCl, 0.5mM spermidine, 0.01% digitonin, 2mM EDTA, 1x Roche cOmplete mini EDTA-free protease inhibitor). Secondary antibody (dilution 1:100) was bound in 25µL per sample of Wash150 Buffer (20mM HEPES-KOH pH7.5, 150mM NaCl, 0.5mM spermidine, 1x Roche cOmplete mini EDTA-free protease inhibitor) for 30min at room temperature. pAG-Tn5 pre-loaded adapter complexes (Epicypher) were added to the nuclei-bound beads for 1hr at room temperature in 25µL of Wash300 Buffer (20mM HEPES-KOH pH7.5, 300mM NaCl, 0.5mM spermidine, 1x Roche cOmplete mini EDTA-free protease inhibitor), then beads were washed and resuspended in Tagmentation Buffer (Wash300 Buffer + 10mM MgCl2) and incubated at 37°C for 1hr in a thermocycler with heated lid. Tagmentation was stopped by addition of EDTA, SDS, and proteinase K. DNA was extracted by Phenol-Chloroform and amplified by PCR using CUTANA High Fidelity 2x PCR Master Mix (EpiCypher) and cycling conditions: 5min at 58°C; 5min at 72°C; 45sec at 98°C; 14 cycles of 15sec at 98°C, 10sec at 60°C; 1min at 72°C. PCR products were cleaned up using SPRI beads (Agencourt) at a ratio of 1.3:1 according to manufacturer’s instructions.

### CUT&Tag Analysis

CUT&Tag data were aligned to the GRCh38 patch 13 human genome e following the Benchtop CUT&Tag v3 protocol (Kaya-Okur et al., 2019). Subsequent to alignment we calculated 1x genome coverage normalization with read centering and read extension using deepTools’ bamCoverage (Ramírez, 2016) then mapped the resulting coverage tracks to regions of interest using bedtools’ map function (Aaron R. Quinlan, 2010). Coverage graphs were plotted using ggplot2 from the tidyverse package in R (Wickham H & E, 2019).

### Antibodies

Antibodies against STAT1 [EPR4407] (ab109320, RRID:AB_10863383), STAT1 [EPR21057-141] (ab234400), STAT1 [EPR23049-111] (ab239360), pSTAT1 Y701 [M135] (ab29045, RRID:AB_778096), pSTAT1 S727 [EPR3146] (ab109461, RRID:AB_10863745), YBX1 [EP2708Y] (ab76149, RRID:AB_2219276), PABPC1 (ab21060, RRID:AB_777008), hnRNPM [EPR13509(B)] (ab177957, RRID:AB_2820246), and PRKDC [Y393] (ab32566, RRID:AB_731981) were purchased from Abcam. Antibodies against FLAG tag [M2] (F1804, RRID: AB_262044) and [M2] (F3145, RRID:AB_259529) were purchased from Sigma Aldrich. Antibodies against DDX3X [D19B4] (8192S, RRID:AB_10860416), hnRNPK [R332] (4675S, RRID:AB_10622190), TRIM28 [C42G12] (4124S, RRID:AB_2209886), PPP2R1A [81G5] (2041S), NCL [D4C7O] (14574, RRID:AB_2798519), MYC tag [71D10] (2278, RRID:AB_490778), IDO1 [D5J4E] (86630, RRID:AB_2636818), pSTAT1 Y701 [58D6] (9167, RRID:AB_561284), and Phospho-Rbp1 CTD (Ser5) [D9N5I] (13523, RRID:AB_2798246) were purchased from Cell Signaling Technology. Antibodies against hnRNPU (14599-1-AP, RRID:AB_2248577) and CDK4 (11026-1-AP, RRID:AB_2078702) were purchased from ProteinTech. The antibody against DUX4 (P2G4) was described previously (Geng et al., 2011). The Goat anti-Rabbit IgG HRP (ThermoFisher #A27036, RRID:AB_2536099) and Rat anti-Mouse IgG HRP for IP (Abcam #ab131368, RRID:AB_2895114) were used as secondary antibodies for western blotting.

### Primer Sequences

#### ChIP-qPCR

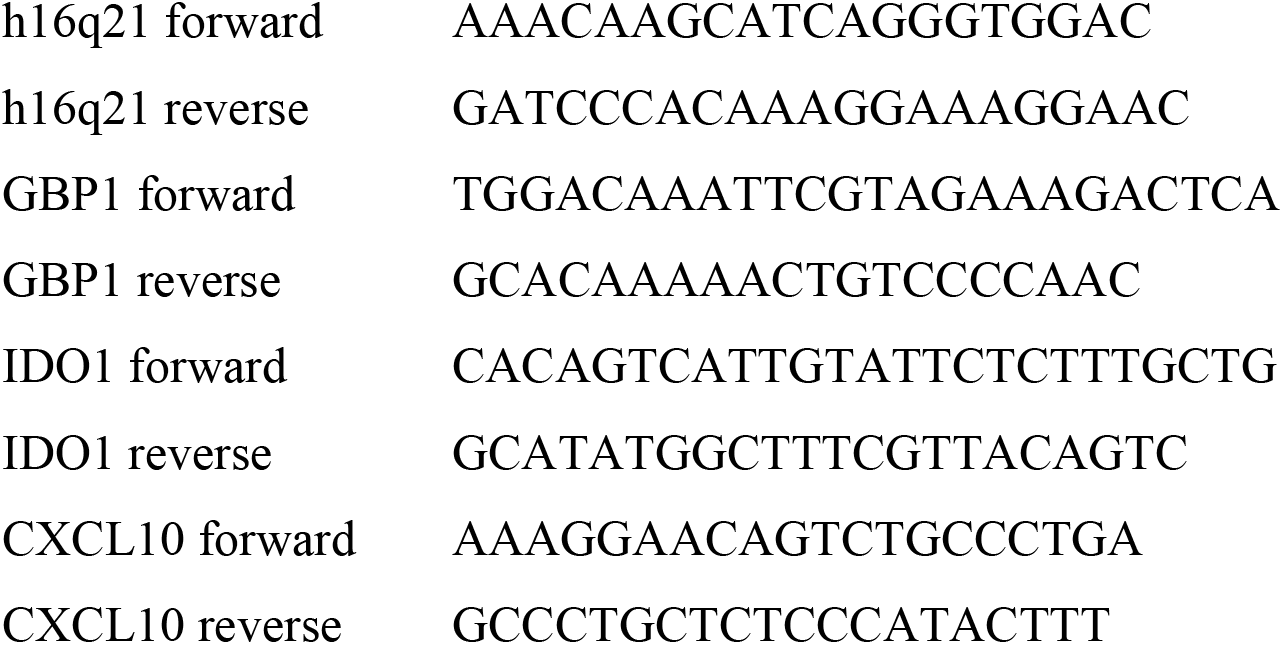

#### RT-qPCR

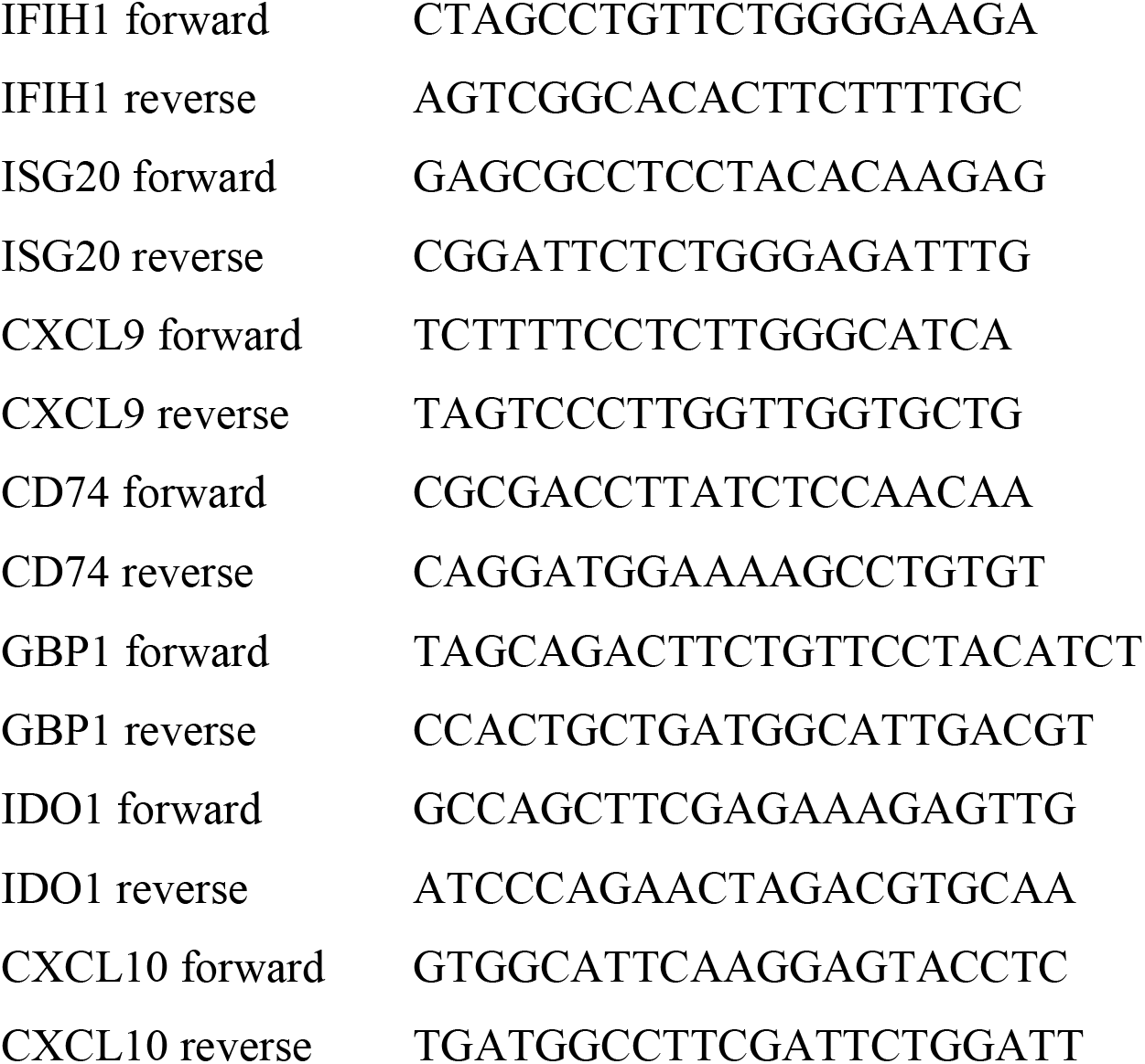

## ACKNOWLEDGEMENTS

We thank Daniel Stetson and Michael Gale for providing advice and reagents. The FHCRC Genomics and Proteomics Cores for outstanding services. Funding was provided by NIH NIAMS AR045203 (SJT), NCI P30 CA015704 Supplement (SJT), T32 HG000035 (AES), The Friends of FSH Research and the Chris Carrino Foundation for FSHD (AES, NAS, SRB, AEC, SJT). We thank Dr. H. Otani and Osaka University for providing the Kitra-SRS cell line.

## AUTHOR CONTRIBUTIONS

Conceptualization (AES, NAS, AEC, SJT); Methodology, Validation, and Investigation (AES, NAS, AEC, SJT); Data Curation (SRB); Writing (AES, NAS, SJT); Supervision and Project Administration (SJT); Funding Acquisition (AES, SJT).

## Materials Availability

Plasmids used in this study will be deposited with Addgene or are available through request to the corresponding author.

## Data availability

RNA sequencing data and CUT&Tag data are available through GEO GSE186244 and GSE209785, respectively. The mass spectrometry proteomics data have been deposited to the ProteomeXchange Consortium via the PRIDE partner repository with the dataset identifier PXD029215. Standard packages were used for RNA sequencing and CUT&Tag analyses (see “Materials & Methods”); Additional code for data processing is available on figshare: https://figshare.com/s/73ab926975aefe2db577

## SUPPLEMENTAL FIGURES

**Supplemental Figure S1.**
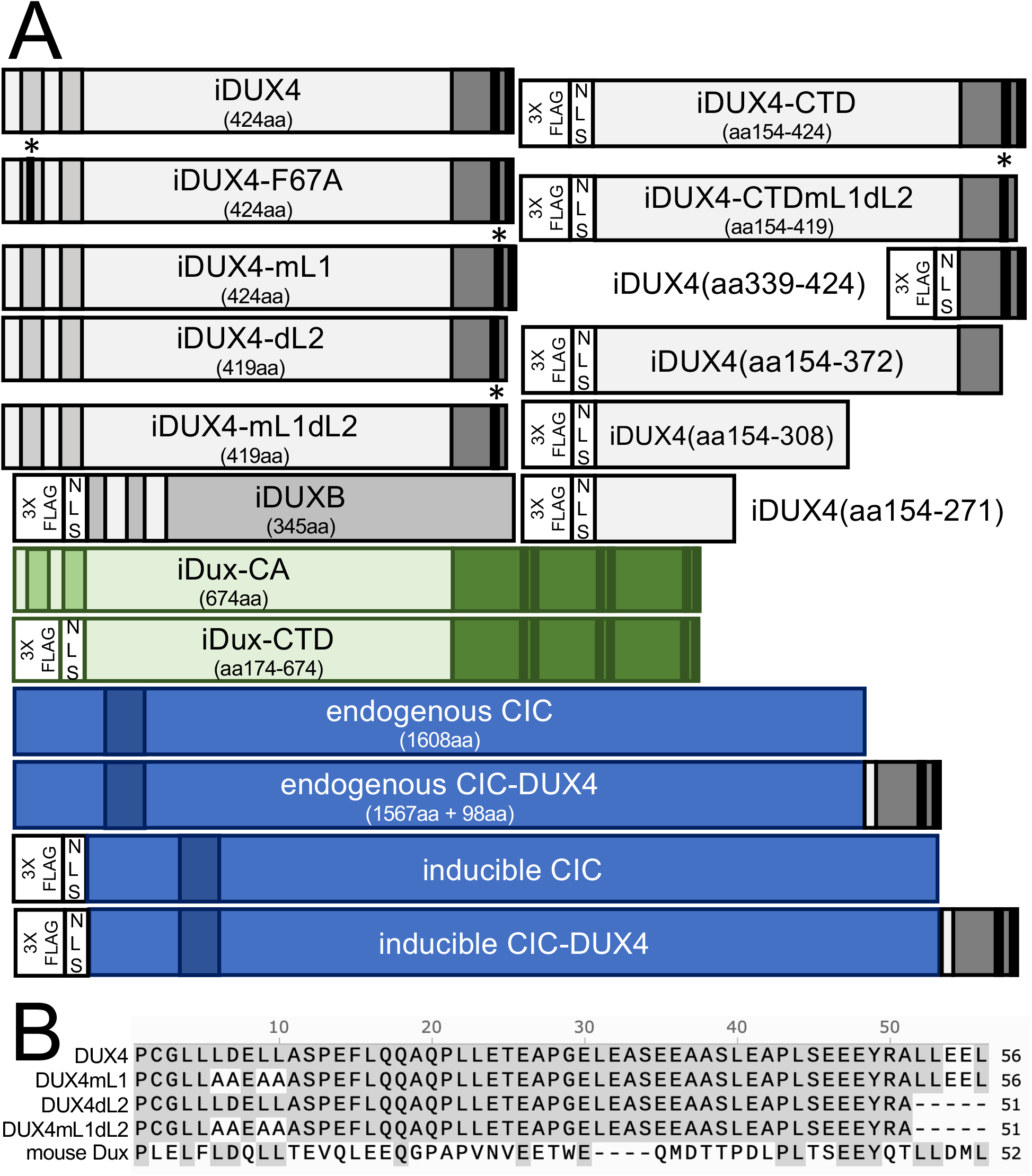
Transgene diagrams. (**A**) Schematic depiction of each transgene used in this study highlighting the N-terminal homeodomains (light grey in DUX4, no fill in DUXB, light green in mDux), DNA-binding HMG box (dark blue in CIC and CIC-DUX4), conserved C-terminal domain (medium grey in DUX4 and CIC-DUX4, medium green in mouse Dux), (L)LxxL(L) (black in DUX4 and CIC-DUX4, dark green in mouse Dux), mutations (* and black bar F67A, * replacement of (L)LxxL(L) with AADEAA), and 3xFLAG-NLS cassette regions (no fill). (**B**) MUSCLE alignment of the terminal ∼50aa of the human DUX4, mutated human DUX4 (mL1, dL2, mL1dL2), and mouse Dux constructs used in this study.

**Supplemental Figure S2.**
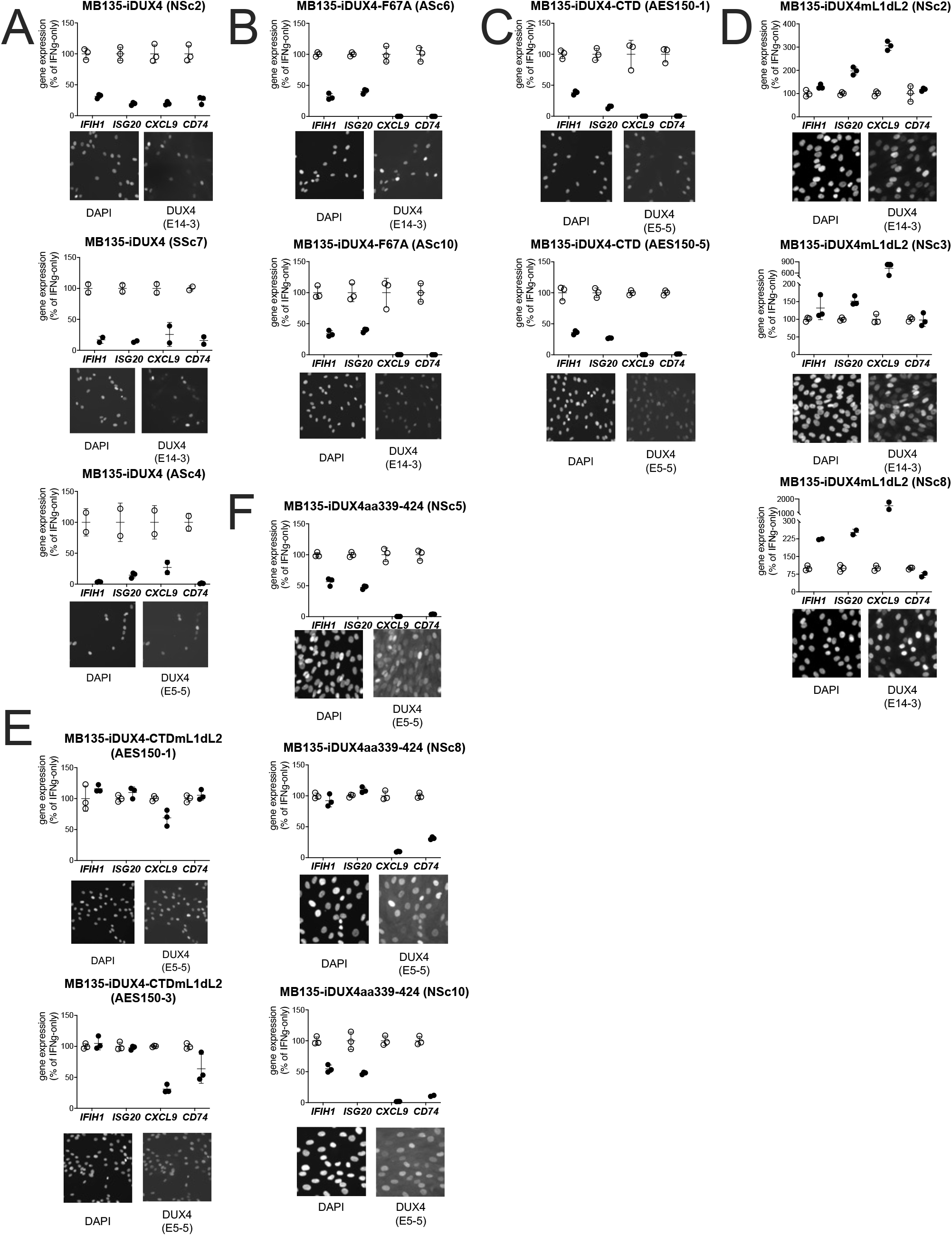
Biological replicates in independent cell lines for each DUX4 construct. Additional subcloned MB135 cell lines of the iDUX4 (**A**), iDUX4-F67A (**B**), iDUX4-CTD (**C**), iDUX4mL1dL2 (**D**), iDUX4-CTDmL1dL2 (**E**), iDUX4aa339-324 (**F**) treated with IFNγ ±doxycycline. RT-qPCR shows ISG expression graphed as a % of IFNγ-only. Data represent the mean ±SD of three biological replicates with three technical replicates each. Immunofluorescence panels show protein expression and nuclear localization using an antibody against the N-terminal (E14-3) or C-terminal (E5-5) residues of DUX4 as appropriate for the construct.

**Supplemental Figure S3.**
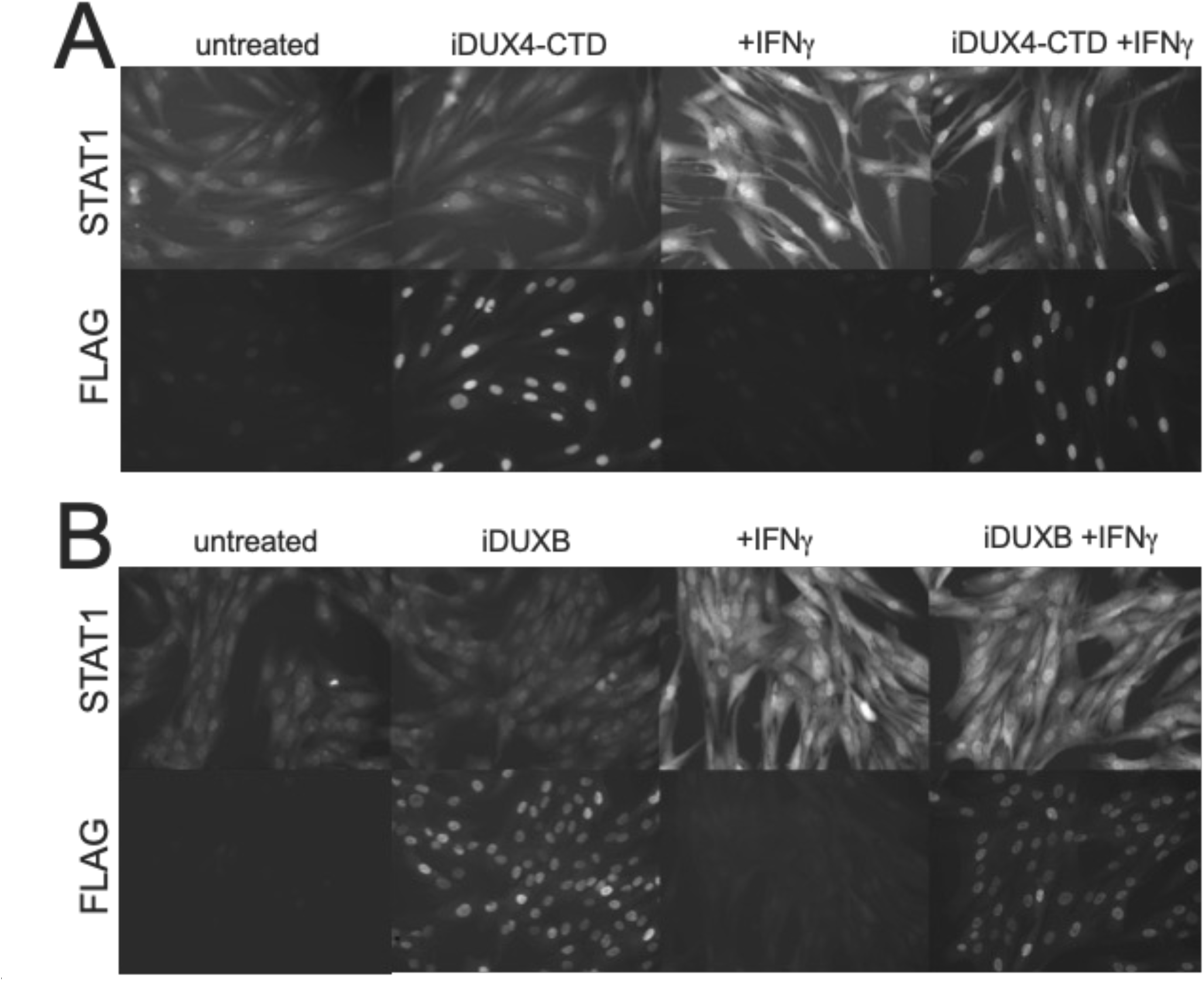
Expression of the DUX4-CTD does not prevent translocation of STAT1 to the nucleus. MB135-iDUX4-CTD (**A**) and MB135-iDUXB (**B**) cells were left untreated, treated with doxycycline, treated with IFNγ, or treated with combination doxycycline/IFNγ, then fixed and stained for STAT1 and either transgene. Both cell lines show good induction and nuclear translocation of STAT1 with IFNγ, with or without doxycycline-induction of the transgene.

**Supplemental Figure S4.**
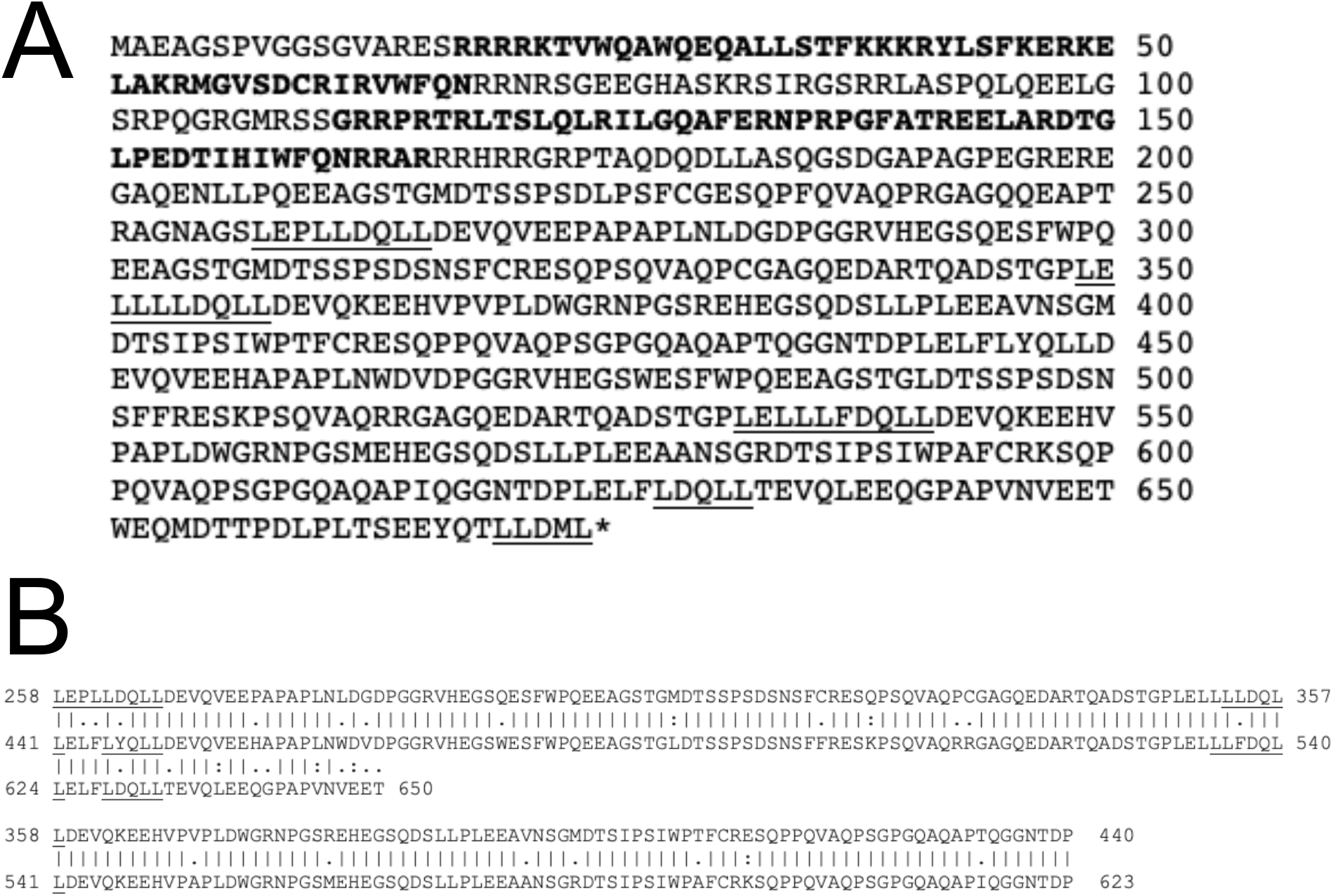
Mouse Dux contains a triplication of the (L)LxxL(L)-containing region. (**A**) Mouse Dux protein sequence with homeodomains in bold and (L)LxxL(L) motifs underlined. (**B**) Alignment of a partial triplication of the mouse Dux protein with aa258-440 aligning with aa441-623 and aa624-650 aligning with aa258-284.

